# Plk1 inhibition delays mitotic entry revealing prophase-specific changes to the phosphoproteome

**DOI:** 10.1101/2024.04.04.588210

**Authors:** Monica Gobran, Luisa Welp, Jasmin Jakobi, Antonio Politi, Henning Urlaub, Peter Lenart

## Abstract

Polo-like kinase 1 (plk1) is a conserved regulator of cell division. During prophase, plk1 phosphorylates direct substrates and is involved in activation of the cyclin-dependent kinase 1 (cdk1). However, the exact functions of plk1 in prophase remain incompletely understood. By testing several cell lines and small-molecule inhibitors, we confirm that plk1 inhibition causes a delay in mitotic entry. We show that cells are delayed in a prophase-like state displaying progressively condensing chromosomes, increased microtubule dynamics, reorganization of the actin cortex, while the nuclear envelope remains intact. We show that during this prolonged prophase cdk1 activity increases gradually over several hours with individual cells stochastically reaching the entry threshold, explaining the highly variable timing of mitotic entry. We then use phosphoproteomics to characterize this prolonged prophase state revealing for the first time phosphosites specific to prophase including several regulators of chromatin organization and the cytoskeleton. Together, we show that plk1 functions as a catalyst of prophase to prometaphase transition, and by using plk1 inhibition as a tool, we identify early changes in the phosphoproteome as the cell prepares for division.

## Introduction

While prophase is unquestionably the first phase of cell division, ‘mitotic entry’ is conventionally defined as the next step; the transition from prophase to prometaphase. The reason behind this definition is that the transition to prometaphase is marked by nuclear envelope breakdown (NEBD), a sudden event easily visible even through a simple microscope. At the molecular level, NEBD coincides with the full activation of cdk1-cyclin B, causing a massive phosphorylation of mitotic substrates, and marking the irreversible commitment to division (Gavet and Pines, 2010; Crncec and Hochegger, 2019). Cells can be easily synchronized in prometaphase by spindle poisons, which enabled the detailed characterization of this state by various methods, including phosphoproteomics (Daub et al., 2008; Dephoure et al., 2008). By stark contrast, the onset, the exact timing, the specific cellular events and the corresponding molecular changes that occur during the preceding prophase state remain poorly characterized, largely due to the lack of protocols for synchronization of cells at this state.

What we do know is that after completion of DNA replication (S-phase), somatic cells enter the second gap phase (G2) to prepare for division. After spending a few hours in G2, cells proceed to prophase. In prophase, chromosomes start to condense, primarily driven by the nuclear condensin II complex (Hirota et al., 2004). The cytoskeleton also starts to reorganize: centrosomes mature, meaning that they recruit additional pericentriolar material (PCM) to increase their microtubule nucleation capacity, and subsequently separate (Lee and Rhee, 2011; Tanenbaum and Medema, 2010). Concomitantly, microtubules begin to change from the interphasic to mitotic dynamics (Ferenz and Wadsworth, 2007). Similarly, the actin cytoskeleton starts to remodel preparing the cell for rounding up. Therefore, stress fibers and focal adhesions disassemble (Chen et al., 2022), and through the activation of the RhoA pathway, cortical contractility is increased (Taubenberger et al., 2020). Furthermore, the disassembly of the nuclear envelope (NE) also begins in prophase with the release of a subset of nuclear envelope proteins (Velez-Aguilera et al., 2020). Due to phosphorylation of proteins of the nuclear pore complex (NPC), nucleo-cytoplasmic transport is also altered (Linder et al., 2017; Nkoula et al., 2023). However, the overall nucleo-cytoplasmic compartmentalization remains intact during prophase.

The above changes in cellular architecture are driven by mitotic kinases, prominently Aurora A and B, polo-like kinase 1 (plk1) and cdk1. For example, in prophase Aurora B phosphorylates histone H3, an early mark of chromosome condensation (Hsu et al., 2000). Plk1 mediates centrosome maturation by phosphorylating the PCM component pericentrin (Lee and Rhee, 2011). Plk1 also contributes to the remodeling of the nuclear envelope through phosphorylation of the nucleoporins Nup98 and Nup53 (Linder et al., 2017; Laurell et al., 2011). In addition, Aurora kinases and plk1 are also key components of the signaling network leading to full activation of cdk1.

In mammalian somatic cells, cdk1-cyclin B complexes are present and ready to be activated already as cells enter G2. However, cdk1 is kept inactive by phosphorylations on Thr14 and Tyr15 by wee1/myt1 kinases (Crncec and Hochegger, 2019). Cdk1 activity then begins to increase gradually in late G2, in a process that remains incompletely understood. A critical event for activation of cdk1 is the dephosphorylation of the Thr14/Tyr15 sites by cdc25 phosphatases. This is a highly regulated process receiving inputs from extracellular (e.g. growth factors) and intracellular (e.g. DNA damage) signaling pathways, and is modulated by multiple auto-activatory feedback loops (Liu et al., 2020). One major pathway identified to activate cdc25 is as follows (Gheghiani et al., 2017): cdk1/2-cyclin A phosphorylates the protein Bora, leading to the activation of Aurora A, which in turn phosphorylates and activates plk1. plk1 then phosphorylates and activates cdc25C, which then dephosphorylates and activates cdk1. Once cdk1 is fully activated, it phosphorylates over a thousand mitotic substrates (Dephoure et al., 2008).

Considering the above, it is clear that the cell begins to reorganize its architecture already in prophase at a time when cdk1 is only partially active. However, we lack a systematic overview of the molecular changes, in particular changes in phosphorylation states, that occur in prophase. Previous approaches used immunolabeling of cells and subsequent FACS sorting (PRIMMUS and its advanced variants), providing rather limited cell numbers of such transient stages as prophase (Ly et al., 2017; Kelly et al., 2021). Another study used highly synchronous cell populations released into mitosis, which relied on very tight 5-minute sampling of cells and focused exclusively on chromatin associated proteins (Samejima et al., 2022). Thus, the inability to obtain a large and homogeneous prophase cell population remains a strong limitation that prevented so far a systematic characterization of prophase by phosphoproteomics.

Here, we investigated the effect of plk1 inhibition to gain insights into cellular events of prophase and their regulation. Previous studies using various cell lines and different methods for inhibiting plk1 showed varied effects ranging from a slight delay to complete inhibition of mitotic entry (e.g. Lénárt et al., 2007; Gheghiani et al., 2017). We confirm this and show that the delay results from a transient or permanent arrest in a prophase-like state. In this prolonged prophase, cells display progressively condensing chromosomes and increased dynamics of the actin and microtubule cytoskeleton, while the nuclear envelope remains largely intact. By using a FRET-based cdk1 activity probe as well as by monitoring the localization of cyclin B, we further show that cdk1 activity increases gradually over several hours during prolonged prophase. As a result, only stochastically and with a substantial delay individual cells reach the cdk1 activity threshold required to progress to NEBD. This establishes plk1 as a catalyst of prophase to prometaphase transition, and explains the highly variable delay of mitotic entry in the absence of plk1 activity. Based on these observations, we then used plk1 inhibition as a tool to enrich cells in prolonged prophase and analyzed them by phosphoproteomics. This provides the first catalog of phosphorylation events that underlie changes in cellular architecture early in cell division.

## Results

### Plk1 inhibition causes delayed mitotic entry with highly variable timing

Previous publications reported somewhat inconsistent effects of plk1 inhibition, therefore we first wanted to compare three cell lines side-by-side that are most commonly used in cell cycle studies: two cancer cell lines, HeLa and U2-OS, and an hTERT immortalized non-cancer cell line, hTERT-RPE1. We treated synchronized populations of cells at release from S-phase arrest. We then filmed the cells stained with the red fluorescent Hoechst derivatives 5’-SiR- or 5’-CP-580-Hoechst (Bucevičius et al., 2019) for 24 hours (Fig. 1A). Mitotic entry (i.e. NEBD) was identified by the collapse of the condensed chromosomes (Fig. 1B). Imaging *per se* had no significant effect on the cells, as DMSO-treated controls progressed though division with normal timing and morphology and consistent with growth rates in culture (Fig. 1B, C). However, the extent of synchrony differed between cell lines with HeLa cells responding best to our single thymidine synchronization regimen, and U2-OS cells being the least synchronous (Fig. 1C).

**Figure 1.**
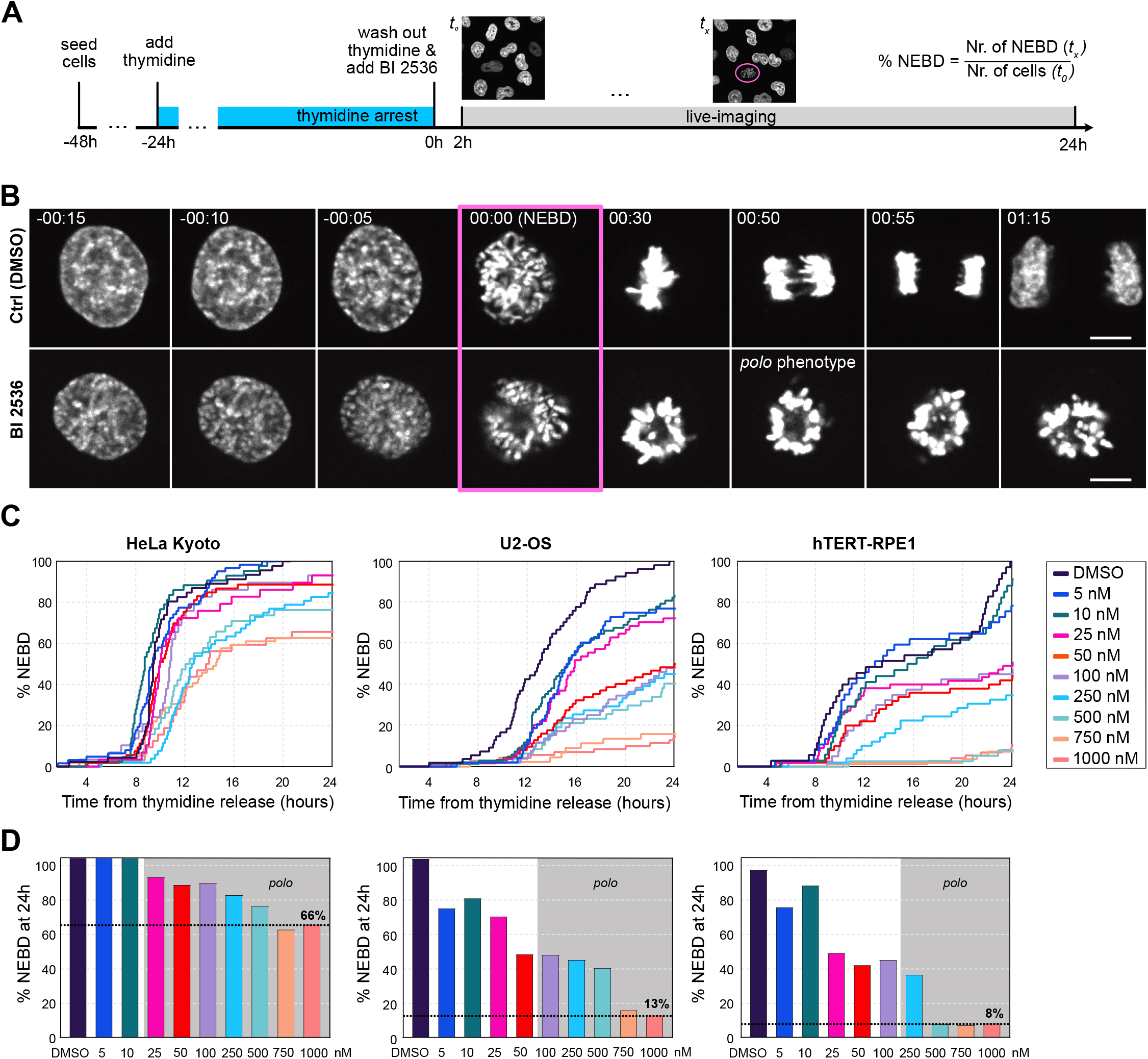
Plk1 inhibition causes delayed mitotic entry with highly variable timing. **(A)** Schematics of the experimental workflow. Images shown in panels below are all cropped and selected frames from such experiments. **(B)** HeLa cells stained with 5-SiR-Hoechst at different stages of mitosis with and without treatment with 250 nM BI 2536. Scale bars are 10 µm. **(C)** Cumulative plots of NEBD timing in response to increasing concentrations of BI 2536 in HeLa Kyoto, U2-OS, and hTERTRPE1 cells. **(D)** Bar charts show the percentage of cells that had undergone NEBD by the end of the experiment, gray areas indicate the concentrations at which the *polo* phenotype appears. For each treatment a total of 30 to 70 cells were analyzed.

We treated cells with increasing concentrations of BI 2536, a well-established and highly specific inhibitor of plk1 (Lénárt et al., 2007; Steegmaier et al., 2007). Starting from low concentrations expected to only partially inhibit plk1, we escalated doses to multiples of which have been demonstrated to suffice for complete inhibition (100 nM for HeLa and U2-OS, 250 nM for hTERT-RPE1 cells (Steegmaier et al., 2007; Petronczki et al., 2007)). As also shown in these and other studies, the fully penetrant *polo* phenotype -- chromosomes arranged in a ring shape -- is a good indicator of complete plk1 inhibition, which we observed above 25, 100 and 250 nM in HeLa, U2-OS and hTERT-RPE1 cells, respectively (Fig. 1B-D). Importantly, we observed the *polo* phenotype to the same penetrance at late time points, arguing against degradation or inactivation of BI 2536 over the 24 h of imaging, consistent with pharmacokinetic data (Steegmaier et al., 2007).

In all cell lines investigated, plk1 inhibition caused a delay in mitotic entry in parallel to an increasing fraction of cells failing to enter mitosis. The extent of the delay and the fraction of the cells that failed to enter mitosis differed substantially between cell lines. In HeLa cells, plk1 inhibition caused an entry delay of 4-5 hours and even at 1 μM (10-times above the full inhibitory dose) two-thirds of the cells entered mitosis. U2-OS cells showed a more extended delay of more than 12 hours and only ∼10% of the cells still entered mitosis at the highest 1 μM concentration. Mitotic entry in hTERT-RPE1 cells was delayed similarly to HeLa, by approximately 4-6 hours, but the proportion of cells entering mitosis was substantially lower, only 8% at higher concentrations (Fig. 1C, D).

Using HeLa cells expressing PCNA-EGFP, we confirmed that the duration of S-phase was unaffected by BI 2536, indicating that the delay occurs in G2/M (Fig. S1A, B). We could also confirm that synchronization had no effect on the delay caused by BI 2536 treatment (Fig. S1C). Furthermore, γ-H2AX staining revealed no significant difference in the number of DNA damage foci, indicating that plk1 inhibition is unlikely to cause this delay by affecting DNA damage (Fig. S1D), consistent with previous results (Gheghiani et al., 2017). To further substantiate our findings, we also tested the comparably selective and potent, but structurally unrelated plk1 inhibitor, GSK 461364 (Gilmartin et al., 2009). Cells showed a very similar delay in mitotic entry as observed with BI 2536, and displayed a fully penetrant *polo* phenotype after NEBD (Fig. S1E, F).

Taken together, our systematic survey across concentrations and cell lines confirms that plk1 activity is required for timely mitotic entry, i.e. NEBD. However, this effect is highly variable not only between cell lines, but also between individual cells in a population. Within a single field of view, we observed cells that enter mitosis almost without delay, other cells are delayed for many hours, or do not enter at all during the observation period. However, in all cell lines investigated, a substantial portion of cells were able to enter mitosis even at the highest inhibitor concentrations indicating that mitotic entry, even if delayed, can occur in the absence of plk1 activity.

### Plk1 inhibited cells are delayed in a prolonged prophase-like state with condensing chromosomes

We next wanted to find out at which stage plk1-inhibited cells are delayed before transitioning to prometaphase and undergoing NEBD. In control cells, closer inspection of the images at high resolution revealed early signs of chromosome condensation, a hallmark of prophase, 10-20 minutes prior to NEBD. Strikingly, in BI 2536-treated cells we observed condensing chromosomes much earlier, several hours preceding NEBD in all three cell lines investigated (Fig. 2A). In order to quantify this effect, we automatically segmented nuclei and used the standard deviation of fluorescence intensities as a measure of chromosome condensation (Neurohr and Gerlich, 2009). By this measure, we were able to detect the start of chromosome condensation in control cells already much earlier, 1-2 hours before NEBD: a slow and gradual increase followed by a steep rise ∼20 min before NEBD (Fig. 2B). Intriguingly, in BI 2536-treated cells we observed chromosome condensation to begin up to 10-15 hours before NEBD. Chromosome condensation then continued up to NEBD, while several cells remained arrested with condensing chromosomes until the end of the recording (Fig. 2B). The timing differed slightly between cell lines (see above), but overall they all showed a similar response (Fig. 2A, B). HeLa cells treated with GSK 461364 also showed an identical phenotype displaying delayed mitotic entry with progressively condensing chromosomes (Fig. S1E-G).

**Figure 2.**
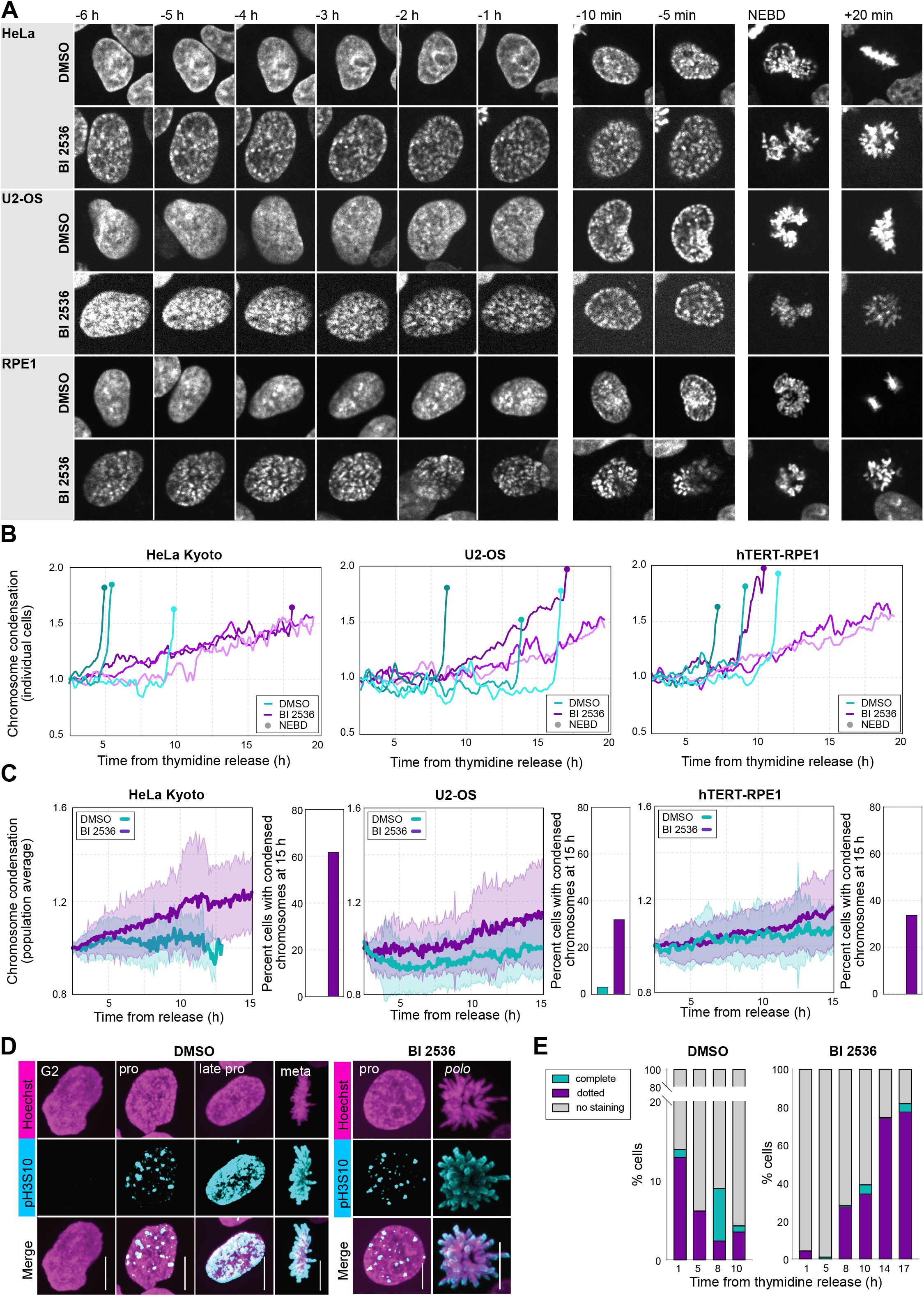
Cells are delayed in a prolonged prophase-like state with condensing chromosomes. **(A)** Montages showing mitotic entry of exemplary control and BI 2536-treated cells stained with 5-SiR-Hoechst. **(B)** Quantification of chromosome condensation using the standard deviation of fluorescence intensity of individual cells on recording similar to shown in (A). **(C)** Population averages of tracked nuclei of measurements as in (B), mean ± standard deviation is shown. **(D)** Immunofluorescence staining of H3S10ph in HeLa Kyoto cells treated with and without 250 nM BI 2536 at different stages of division. **(E)** Quantification of different morphologies on images similar to (D) in cells fixed at different times after thymidine release. The quantification is done only on attached cells. For (A) and(D) scale bars are 10 µm.

The above effects observed in single cells was also reflected on population averages quantified on all tracked cells in a field of view. Control cells showed no systematic trend in chromosome condensation during G2. By contrast, in BI 2536 treated cells, chromosome condensation progressively increased resulting from more and more cells starting to condense chromosomes in the population hours before NEBD (Fig. 2C). The increase in chromosome condensation observed in populations of live cells correlated tightly with a progressive increase in the number of cells showing phospho-histone 3 serine 10 (pH3S10) staining (Fig. 2C-E). At the late time point, 17 hours after thymidine release, up to 80% of the BI 2536-treated cells showed the dotted pH3S10 staining pattern typical of prophase (Fig. 2E) (Hirota et al., 2005).

Taken together, our data show consistently across cell lines and by using two structurally unrelated inhibitors that when plk1 is inhibited, cells are delayed in a prolonged prophase-like state with progressively condensing chromosomes displaying the early, dotted pH3S10 marks. Cells spend several hours in this stage before finally undergoing NEBD, while many of the cells remain permanently blocked in this state until the end of the 24 h observation period.

### Prolonged prophase cells display dynamic microtubules, signs of rounding up and intact nuclear envelope

Next we wanted to see whether delayed cells show hallmarks of prophase other than chromosome condensation. We first visualized microtubules by using low concentrations of the fluorescently labeled taxane, 6-SiR-CTX (Bucevicius et al., 2020). Confirming previous reports, while BI 2536-treatment did not visibly change the total amount of microtubules in prophase and prometaphase cells, microtubule organization was strongly affected (Fig. 3A). As plk1 activity is required to recruit the pericentriolar material to centrosomes, in BI 2536-treated cells centrosomal asters do not form, and the microtubule network shows a de-focused organization (Lénárt et al., 2007).

**Figure 3.**
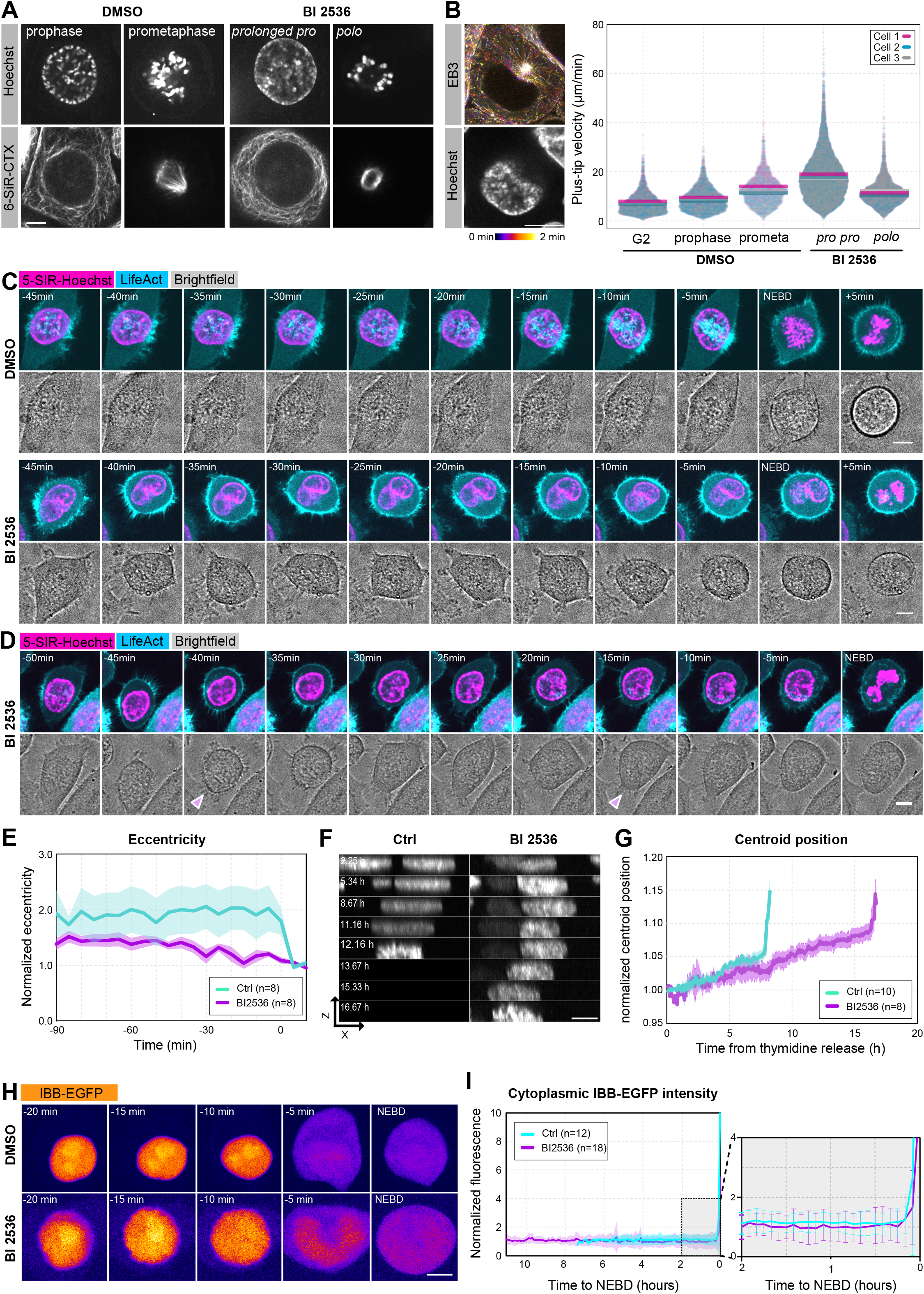
Prolonged prophase cells display dynamic microtubules, signs of rounding up and an intact nuclear envelope. **(A)** HeLa cells stained with 5-580-CP Hoechst and 6-SiR-CTX at different stages of mitosis treated with or without 250 nM BI 2536. **(B)** HeLa cells expressing EB3-mEGFP and stained with 5-SiR-Hoechst (EB3-mEGFP is shown as a time projection of 81 frames every 2 s). Plus tip velocities of all EB3 tracks recorded from 3 cells at the indicated stages of division. Horizontal bars represent the mean for each cell. **(C)** HeLa cells expressing LifeAct-mEGFP and stained with 5-SiR-Hoechst treated with and without BI 2536. **(D)** Images of a BI 2536-treated HeLa cell a few hours before undergoing NEBD. Pink arrows show rounding up events. **(E)** Measurement of the eccentricity of the LifeAct-mEGFP labeled cell outline in experiments similar to (C). Data is normalized to the mean eccentricity of the 2 frames after NEBD. **(F)** XZ ‘side view’ of cells stained with 5-SiR-Hoechst. Times are relative to thymidine release. **(G)** Quantification of data shown in (F) by calculation of the position of the centroid (center of mass) in the Z-axis normalized to the mean position in the first 5 frames. **(H)** Selected frames from a time lapse showing the localization of IBB-mEGFP in the time leading to NEBD in control and BI 2536-treated cells. **(I)** Quantification of IBB-mEGFP mean cytoplasmic fluorescence intensity in control and BI 536-treated cells on recordings similar to (H). Data is normalized to the mean intensity in the first 5 frames of imaging. Inset showing the time window around NEBD is shown on the right. For (A), (C ), (D), (F), and (H): scale bars are 10 µm.

To assay the dynamics of these microtubules, we imaged cells stably expressing EB3-mEGFP at different phases of mitosis staged by chromosome morphology (Fig. 3B). We tracked microtubule tips using the plusTipTracker software to quantify microtubule dynamics (Applegate et al., 2011). In control cells, this analysis revealed dynamics consistent with data published earlier (Ferenz and Wadsworth, 2007), showing an increase in plus-tip velocities as cells progress from G2 to prophase and then to prometaphase (Fig. 3B). Intriguingly, cells delayed in prolonged prophase by BI 2536 showed plus-tip growth velocities significantly higher than prophase, and even higher than prometaphase of DMSO-treated control cells (Fig. 3B). Thus, cells in prolonged prophase not only display condensed chromosomes, but also show increased, mitotic-like microtubule dynamics.

Secondly, we monitored the dynamics of the actin cytoskeleton by imaging cells stably expressing the actin marker, LifeAct-mEGFP (Riedl et al., 2008). This marker visualized a strong cortical staining in BI 2536-treated cells delayed in prolonged prophase, comparable or even stronger than the cortical label in control cells at metaphase (Fig. 3C). Concomitantly, we observed attempts of BI 2536-treated cells to detach and to begin to round up 1-2 hours before NEBD, much earlier than this occurs in control cells (Fig. 3C). We could see cells partially rounding up, and in several instances we observed an oscillatory behavior with cells rounding up and then attaching again during prolonged prophase (Fig. 3D). To quantify these effects, we measured the eccentricity of the cell outlines (marked by LifeAct-EGFP). This showed a slow, continuous decrease in BI 2536-treated cells as opposed to the sudden decrease in eccentricity shortly before NEBD in control cells (Fig. 3E). The change in cell shape was also mirrored by a change in nuclear shape visible on z-sections and reflected by a continuous increase of the position of the centroid of the nucleus along the z-axis (Fig. 3F, G).

Next, we monitored the state of the nuclear envelope (NE) in cells delayed in prolonged prophase. To this end, we stably expressed the nuclear import substrate IBB-EGFP, the importin-β binding domain of importin-α fused to EGFP (Dultz et al., 2008). Confirming earlier reports, in DMSO-treated control cells, IBB-EGFP started to be released from the nucleus ∼5 min before NEBD (Fig. 3H). BI 2536-treated cells showed no noticeable difference to controls by visual inspection, and as quantified by measuring the increase in IBB-EGFP intensity in the cytoplasm (Fig. 4I). Consistently, we did not observe a difference in the release kinetics of two other NE markers, the inner nuclear membrane protein LBR, and the nucleoporin Nup107 (Fig. S2A-C).

**Figure 4.**
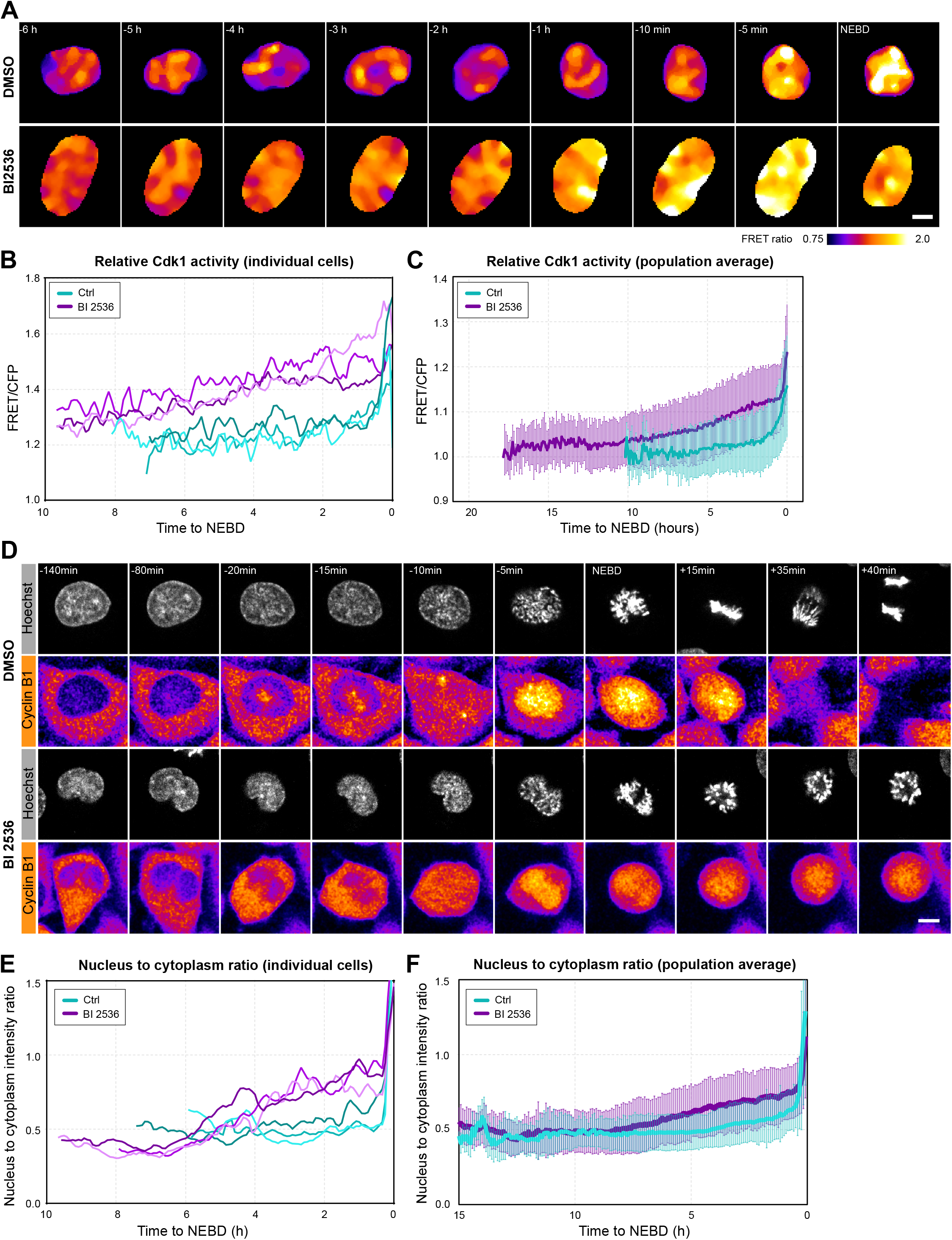
Cdk1 activity increases gradually over hours during prolonged prophase. **(A)** Ratio images (YFP/CFP emission at CFP excitation) in a stable HeLa cell line expressing the Eevee-spCDK FRET probe. Selected frames showing cells entering mitosis. Time is relative to NEBD. **(B)** Quantification of FRET ratio changes of the Eevee-spCDK FRET probe in individual control and BI 2536-treated cells. **(C)** Quantifications as in (B) showing averages in control and treated populations of cells. **(D)** Live images of exemplary control and BI-2536 treated HeLa cells stably expressing Cyclin B1-mVenus while entering mitosis. Time is relative to NEBD. **(E)** Quantification of Cyclin B1 nucleus to cytoplasm intensity ratio in individual control and BI 2536-treated cells. **(F)** Quantifications as in (E) showing averages in control and treated populations of cells. For (A) and (D): scale bars are 10 µm.

Together, these data indicate that mitotic reorganization of the actin cortex as well as the microtubule cytoskeleton begins during prolonged prophase, several hours before NEBD. It is intriguing that both microtubule dynamics and cortical actin appear to be over-activated to levels even beyond metaphase of control cells. This suggests that the regulatory kinases may be more active, or that due to the extended time spent in prophase, substrates may reach higher levels of phosphorylation. However, while a few of its components may be phosphorylated and partially released, we show that the nuclear envelope remains intact and transport competent during prolonged prophase.

### During prolonged prophase cdk1 activity increases gradually

Thus, prolonged prophase cells show clear signs of mitotic reorganization: chromosome condensation and increased dynamics of the actin and microtubule cytoskeleton. In normal prophase, these changes coincide with activation of cdk1 and are largely mediated by cdk1 activity. Cdk1 activation is also coupled to nuclear accumulation of cdk1-cyclin B1 during prophase. Nuclear accumulation of cdk1-cyclin B1 has been shown to be critical for complete activation of cdk1, and to be required for driving the irreversible transition to prometaphase (Santos et al., 2012; Gavet and Pines, 2010).

To visualize changes in cdk1 activity during prolonged prophase, we established a stable cell line expressing an optimized, Förster Resonance Energy Transfer (FRET) based cdk activity biosensor, Eevee-spCdk (Sugiyama et al., 2023). In control cells, the sensor showed no change during G2 phase followed by a rapid increase in FRET starting ∼20 min before NEBD (Fig. 4A-C), consistent with previous observations made using a similar FRET probe (Gavet and Pines, 2010). This increase in FRET was completely abolished by the cdk1 inhibitor RO 3306 (Fig S3C). Intriguingly, in BI 2536-treated cells we observed a much slower, continuous and gradual increase in cdk activity over several hours during prolonged prophase (Fig 4A, B). In some cells this slow increase transitioned into a steeper slope shortly followed by NEBD in those cells. We observed consistent kinetics in cell populations as well (Fig. 4C).

To visualize changes in the localization of cdk1-cyclin B1, we imaged HeLa cells expressing endogenously tagged cyclin B1-mVenus. In control cells, we could confirm the previously observed nuclear re-localization of cyclin B1 (Gavet and Pines, 2010), the kinetics of which closely matched the activation of cdk1 starting ∼20 min before NEBD (Fig. 4D-F). Strikingly, in BI 2536 treated cells the slow and gradual increase in cdk activity detected by the FRET probe was almost identically mirrored by slow and gradual relocalization of cyclin B1 to the nucleus. We quantified this kinetics in individual cells as well as on cell populations revealing a slow increase occurring over several hours and eventually reaching similar levels to control prometaphase cells at NEBD (Fig. 4D-F). To further confirm these results, we also stained the endogenous cyclin B1 protein by immunofluorescence in an untagged HeLa cell line and used chromosome morphology to stage cells. In the control sample we found cyclin B1 to localize to the cytoplasm in cells with uncondensed chromosomes (G2), while in cells with condensed chromosomes (prophase), cyclin B1 was enriched in the nucleus (Fig. S3A, B). In the BI 2536-treated sample, we observed cytoplasmic localization in cells with non-condensed chromosomes, similar to controls. Different to controls, in cells with condensed chromosomes, we observed an equilibration of cyclin B1 between the cytoplasm and nucleus, with preferential nuclear localization in cells arrested in prolonged prophase for longer periods (17 h) (Fig. S3A, B).

Taken together, the cdk activity FRET probe and cyclin B1 localization consistently evidence a slow and gradual activation of cdk1-cyclin B1 occurring over several hours during prolonged prophase. Therefore, cdk activity reaches the threshold required for NEBD only after a long delay and stochastically in individual cells. Many cells never reach this threshold and remain arrested in prolonged prophase. This explains the delayed and highly variable timing of mitotic entry, and establishes plk1 as a catalyst of prometaphase transition.

### Phosphoproteomic analysis of the prolonged prophase state

Thus, our data combined demonstrates that plk1 inhibition arrests cells for several hours in a pro-phase-like state. This offers an opportunity, for the first time, to collect a large homogenous population of prophase cells and characterize this state by phosphoproteomics.

Therefore, we designed an experiment to quantitatively compare the following states (Fig. 5A): (i) attached cells in G2 5 hours after release from a 24 hour thymidine arrest; (ii) shaken-off prometaphase cells arrested by the kinesin-5 inhibitor, S-trityl-L-cysteine (STLC) (Kaan et al., 2011) 13.5 hours after thymidine release; and (iii) cells treated with 250 nM BI 2536 collected 13.5 h after thymidine release. This latter sample was further divided into two: after a gentle shake-off, suspended cells were collected separately from attached cells, resulting in *polo* and prolonged prophase cell populations, respectively. As confirmed by Hoechst staining, the floating cells were almost exclusively (>98%) cells arrested in prometaphase showing the *polo* phenotype (the others being mostly dead cells), while the vast majority (>85%) of attached cells were in prolonged prophase (the other 15% being mostly cells in G2). In this setting, we used the G2 sample as reference relative to which we were able to detect mitotic phosphorylations in the STLC-arrested prometaphase state. BI 2536-treated *polo* prometaphase samples are then expected to display mitotic phosphorylations without plk1-dependent sites, and the attached, prolonged prophase cells to show prophase specific phosphosites, also without plk1-dependent sites.

**Figure 5.**
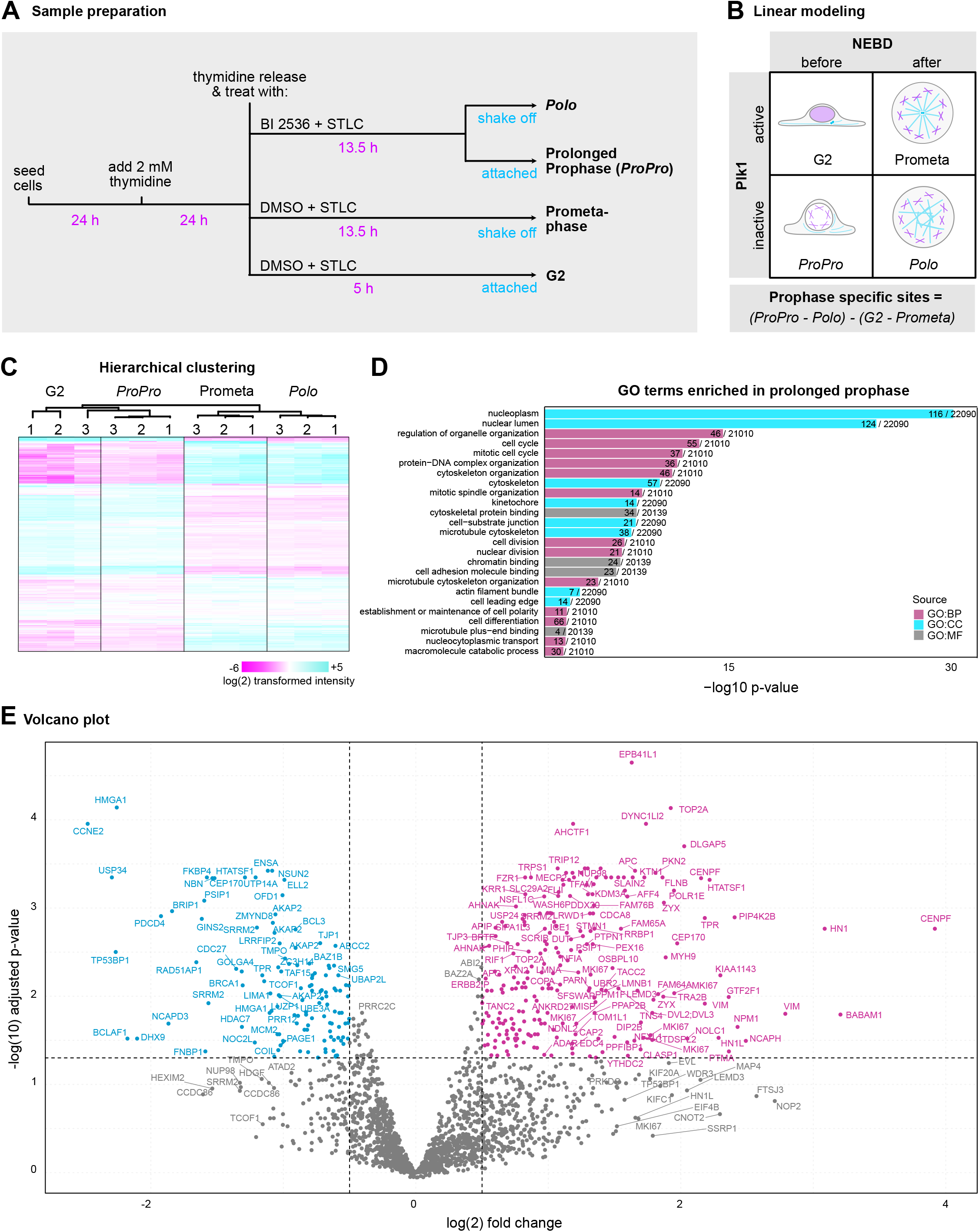
Phosphoproteomic profiling of prolonged prophase. **(A)** Experimental design of the proteomics experiment. Each sample had 3 biological replicates. **(B)** Scheme showing the 2-by-2 factorial design upon which linear modeling was used for the analysis of the dataset. **(C)** Hierarchical clustering of the normalized log(2) transformed intensities of the phosphosites from 4 samples and 3 replicates (denoted by 1, 2, 3). **(D)** Gene ontology terms enriched in the significantly hyperphosphorylated prolonged prophase specific peptides after applying linear modeling. **(E)** Volcano plot showing the differentially phosphorylated prolonged prophase specific peptides after applying linear modeling. The log(2) fold change threshold is set at 0.5 below and above 0 and the significance threshold is set at 0.05 adjusted p-value.

We prepared three biological replicates and used 6-plex Tandem Mass Tag (TMT) isobaric labeling to quantify differences. For analysis of this dataset, we employed linear modeling with a 2-by-2 factorial design using +/- BI 2536 treatment and mitotic time (pre- or post-NEBD) as the two axes. This analysis enabled us to derive the significant differences specific to prolonged prophase (Fig. 5B).

The phosphoproteomics experiment identified 14,994 phosphosites, of which, after filtering, selection, transformation and imputation of the dataset, 2,164 phosphosites could be analyzed (Table S1 and S2). The proteomics experiment identified 7,793 protein groups, of which 5,479 could be analyzed. We did not detect a significant change in the abundance of the phosphoproteins, which were analyzed further. Phosphosite intensities were then normalized showing an overall similar distribution of intensities across biological replicates (Fig S4A).

Hierarchical clustering of the phosphoproteomes revealed a strong separation between pre-NEBD (G2 and prolonged prophase) and post-NEBD (prometaphase and *polo*) samples (Fig. 5C). Within these categories, replicates (G2 vs. prolonged prophase, prometa vs. *polo*) formed distinct clusters. This confirms, firstly, that the variability between replicates is substantially smaller than the difference between treatments. Secondly, the large difference between prolonged prophase and *polo* samples additionally validates our approach to separate suspended *polo* and attached prolonged prophase cells.

Prometa and *polo* samples showed the typical signature of massive mitotic phosphorylation. When comparing the two, the main differences between them were de-phosphorylations on several known targets of plk1 in the *polo* sample (Fig. S4B). Consistently, analysis of the differentially phosphorylated sites revealed plk1 as the main kinase predicted to phosphorylate these sites (Fig. S4C). This validates the specific inhibition of plk1 by BI 2536. By contrast, G2 and prolonged prophase samples lacked most mitotic phosphorylations, making prolonged prophase overall much more similar to the G2 than the M-phase samples (Fig. 5C).

Gene ontology (GO) analysis identified terms enriched in prolonged prophase related to nuclear organization, cytoskeleton and the cell cycle -- an overall signature consistent with the phenotypic changes we observed in prolonged prophase (Fig. 5D). Of the 389 peptides differentially phosphorylated in prolonged prophase (and passing significance thresholds), 246 were phosphorylated and 143 were de-phosphorylated (Fig. 5E, Table S3). Kinase enrichment analysis of hyperphosphorylated peptides in prolonged prophase identified the CMGC kinase family, containing many of the pivotal cell cycle kinases, including cdk1 (Fig. S4D).

Thus, by using BI 2536 to delay cells, we were able to harvest a highly enriched population of cells in prolonged prophase for phosphoproteomic analysis. By combining quantitative analysis of these and samples at other stages of mitosis, we were able to derive the phosphosites that are specifically phosphorylated or dephosphorylated in prolonged prophase. This analysis revealed that prophase is a state very distinct from prometaphase, featuring a limited and specific set of phosphosites, while lacking most mitotic phosphorylations.

### Proteins phosphorylated in prolonged prophase are consistent with cellular phenotypes

Next, we carefully went through the list of proteins differentially phosphorylated in prolonged prophase screening for candidates with functions related to the observed phenotypes. One of the most prominent changes in cellular organization we observed during prolonged prophase is the condensation of chromosomes. Indeed, we detected changes in phosphorylation on two different condensin subunits, NCAPH and NCAPD3 (Fig. 6A). Interestingly, these two subunits belong to two different complexes, condensin I and II, which localize differently to cytoplasm and nucleus, respectively (Hirota et al., 2004). Additionally, we detected changes on two phosphosites of topoisomerase-2α, a DNA de-catenating enzyme with well-established role in chromosome condensation (Nielsen et al., 2020). We also found one phosphosite on CDCA2/Repo-man (Vagnarelli et al., 2006), and prominently, we identified five differentially phosphorylated peptides in KI-67, a protein that coats the surface of chromosomes and is required for their individualization (Cuylen et al., 2016). Additionally, we detected phosphopeptides in the centromere/kinetochore components CDCA8/Borealin and INCENP, two components of the chromosome passenger complex (Jelluma et al., 2008; Xu et al., 2009), and three phosphorylations on CENP-F, which is essential for kinetochore assembly (Vergnolle and Taylor, 2007). We also identified phosphorylation of the TTK/MPS1 kinase, which regulates the assembly of the mitotic checkpoint complex on the kinetochore (Schweizer et al., 2013). Furthermore, we observed phosphorylation of the SWI/SNF and ISWI chromatin remodeling complexes, and histone modifying enzymes, which have been proposed to be involved in chromosome condensation (Wilkins et al., 2014). Taken together, our analysis reveals a set of proteins that are differentially phosphorylated in prophase with established or predicted roles in chromatin organization, chromosome condensation and individualization, as well as in centromere/kinetochore assembly.

**Figure 6.**
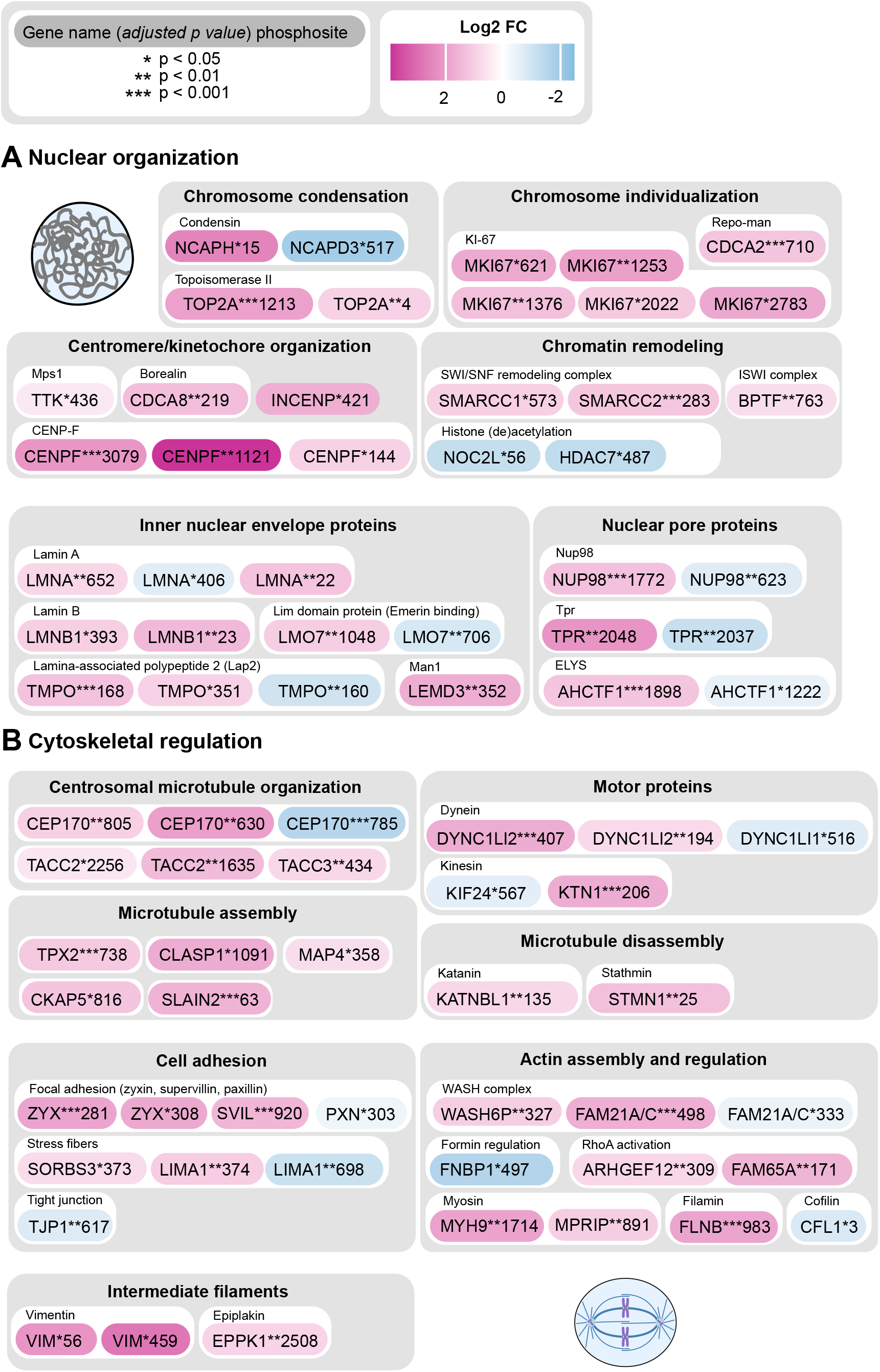
Detailed analysis of the proteins phosphorylated in prolonged prophase. Differentially phosphorylated peptides specific to prolonged prophase after applying linear modeling grouped according to their cellular functions. The gene names of the corresponding peptides are shown, the asterisks mark the adjusted p-value with ***, **, * corresponding to p ≤ 0.001, 0.01, and 0.05, respectively, followed by the position of the phosphosite within the amino acid sequence of the protein. The color coding corresponds to the log(2) fold change of intensity values.

More generally affecting nuclear organization, we also identified several nuclear envelope components to be differentially phosphorylated in prolonged prophase (Fig. 6A). This includes the lamins, lamin A and lamin B. We also identified additional proteins of the inner nuclear membrane, including the lamina-associated polypeptide 2 (LAP2, TMPO), Man1/LEMD3 and the Lim-domain protein (LMO7), a binding partner of Emerin. We found differential phosphorylations on proteins of the nuclear pore complex (NPC), specifically on ELYS, TPR, and Nup98. These changes indicate that besides the lamina, the associated protein network at the inner nuclear envelope as well as the nuclear pores begin to remodel during prophase. However, these changes are specific and restricted to a few sites as compared to the massive phosphorylation of these proteins in prometaphase. This is well consistent with the limited changes observed in nuclear envelope organization and nuclear transport during prophase, while the overall structure of the nuclear envelope remains intact at this stage (Dultz et al., 2008; Lénárt and Ellenberg, 2006). A complete disassembly of the nuclear envelope, nuclear pores and complete depolymerization of the lamina occurs at NEBD, as cells transition to prometaphase and cdk1 is fully activated (Beaudouin et al., 2002). Intriguingly, Samejima and coworkers also found LAP2 and Man1 to be released early from chromatin in prophase in their analysis of chromatin associated proteins (Samejima et al., 2022). Furthermore, our data is also consistent with earlier work showing that phosphorylation of Nup98 (along with Nup53) is an early step in NPC disassembly during prophase (Linder et al., 2017).

Besides nuclear organization, changes in cytoskeletal dynamics are most prominent during prolonged prophase. Consistently, we identified many regulators of the microtubule cytoskeleton to be differentially phosphorylated (Fig. 6B). We identified phosphorylations on the centrosomal protein CEP170 (Conduit et al., 2015), as well as the centrosome associated microtubule regulator TACC2 (Peset and Vernos, 2008). Intriguingly, on the spindle assembly factor TPX2 we detected a phosphorylation on S738. This site has been identified and functionally characterized earlier as a critical ‘early riser’ prophase phosphorylation event (Ly et al., 2017). We additionally identified other regulators of microtubule assembly including CLASP1 and MAP4 (Samora et al., 2011), as well as CKAP5 (Ali et al., 2023) and SLAIN2 (van der Vaart et al., 2011). We also found differential phosphorylations on the microtubule disassembly factors Katanin and Stathmin (Cassimeris, 2002), which may function to sever long interphase microtubules. In addition, we identified differential phosphopeptides in motor proteins including regulators of kinesins and the intermediate light chain 2 of cytoplasmic dynein (DYNC1LI2), known to be involved in centrosome separation (Raaijmakers et al., 2012).

We also identified several regulators of the actin cytoskeleton (Fig. 6B). We observed differential phosphorylations in the focal adhesion components zyxin, supervillin and paxillin, and in a regulator of stress fibers, LIMA1 (Chen et al., 2022). We also detected phospho-changes on components of tight junctions as well as intermediate filaments. Additionally, we detected changes indicative of activation of the Rho pathway, the main pathway responsible for increasing cortical contractility and tension as cells round up (Taubenberger et al., 2020). These proteins include the myosin phosphatase Rho interacting protein (MPRIP), the Rho guanine nucleotide exchange factor 12 (ARHGEF12) and the Rho family-interacting cell polarization regulator 1 (FAM65A/RIPOR1). Somewhat surprisingly, we also identified phosphorylation on several subunits of the WASH complex (FAM21A, FAM21C, WASH6P), an activator of the Arp2/3 actin nucleation complex (Rottner et al., 2010). Intriguingly, WASH has been recently shown to be involved in nucleation of actin filaments at centrosomes (Farina et al., 2019), but whether this has a relevance in prophase remains unknown.

Together, the detected changes on cytoskeletal regulators are well consistent with the disassembly of interphase microtubules and actin structures (focal adhesions, stress fibers), and the concomitant increase in microtubule dynamics and activation of the Rho pathway preparing the cells for rounding up. In summary, we identified changes in the phosphorylation of many proteins with a likely role in the early reorganization of cellular architecture during prophase. In addition to the 53 proteins shown on Figure 6, a complete list of all proteins (corresponding to 389 peptides) modified in prophase, are listed in Table S3 - a remarkably short and specific list of proteins compared to the massive phosphorylation taking place in prometaphase (1859 modified peptides).

## Discussion

The role of plk1 in the regulation of mitotic entry has been studied earlier, but the specific stage at which plk1 inhibition delays cells during mitotic entry, and generally whether plk1 is essential for mitotic entry remained somewhat contentious. To find answers to these questions, we inhibited plk1 in three different cell lines using a wide range of concentrations of the highly-specific small-molecule inhibitors BI 2536 and GSK 461364. By using automated live-cell imaging, we could confirm that plk1 is critical for timely mitotic entry in all cell lines investigated. However, the effect is highly variable between cell lines as well as between individual cells: it ranges from a slight delay of a couple hours to a complete block of mitotic entry - consistent with variable results reported previously. By analyzing cellular phenotypes in detail, we could show that plk1 inhibition delays cells in a prophase-like state. Cells in this *prolonged prophase* display defining hallmarks of prophase: progressively condensing chromosomes, increased microtubule dynamics and remodeling of the actin cytoskeleton preparing cells for rounding up. By contrast, the nuclear envelope remains largely intact.

We show that during prolonged prophase cdk1 activity increases slowly over several hours. After this long delay, stochastically a few cells do eventually reach the cdk1 activity threshold required to undergo NEBD and transition to prometaphase. However, many cells never reach this threshold and remain permanently arrested in prolonged prophase. We thus conclude that plk1 acts as an effective catalyst of prometaphase transition, ensuring that NEBD occurs robustly and within a narrow time window. This is consistent and is explained by earlier work demonstrating that plk1 is an important component of the pathway leading to activation of cdk1 through cdc25C1 (Gheghiani et al., 2017). In the absence of plk1 activity, cells are arrested transiently or permanently in prolonged prophase and NEBD occurs with a substantial delay or is blocked completely. However, our data show that albeit with substantial delay and low efficiency, cells are principally able to enter mitosis in the absence of plk1 activity.

While in many of its aspects prolonged prophase recapitulates normal prophase, it is important to note that due to the lack of plk1 activity, it also differs from it. Firstly, in an unperturbed prophase, plk1 mediates centrosome maturation by phosphorylating pericentrin and other centrosomal proteins (Lee and Rhee, 2011). Secondly, plk1 also phosphorylates the nucleoporins Nup98 and Nup53 facilitating the permeabilization of the nuclear envelope in prophase (Linder et al., 2017). We hypothesize that the lack of these plk1-mediated phosphorylations may actually be critical to keep the nuclear envelope intact and transport competent during prolonged prophase. In normal prophase, these early changes to the nuclear envelope have been proposed to facilitate nuclear accumulation of cdk1-cyclin B (Gavet and Pines, 2010), which is then key to full activation of cdk1, driving NEBD and transition to prometaphase (Santos et al., 2012). Plk1 inhibition may prevent these changes in nuclear transport causing the observed slow and limited relocalization of cyclin B to the nucleus and thus slow and gradual increase in cdk1 activity.

It is important to keep these differences in mind, but in other aspects, cellular phenotypes of prolonged prophase closely mirror an unperturbed prophase. Prolonged prophase thus serves as a model for an otherwise inaccessible transient stage of mitosis, early, before cdk1 is fully activated. As detailed in the Results section, our hit list identified, among several others, differential phosphorylation of many regulators of nuclear organization and the cytoskeleton well-consistent with observed cellular phenotypes. This provides the first comprehensive, while likely still incomplete, list of molecular changes that occur during prophase.

With regard to cell cycle regulation, it is intriguing to consider the model recently published by the Hochegger and Novak groups (Rata et al., 2018). They propose that mitotic entry relies on two interlinked bistable switches, the positive feedback loop activating cdk1, and the feedback through Greatwall leading to the inactivation of the PP2A:B55 phosphatase. Their model implementing these two switches predicts an intermediate but normally hidden steady state between G2 and prometaphase - the prophase state. The model also predicts that this hidden state can be stabilized by ‘disabling’ one of the switches. The authors validate this prediction by showing that by acute inhibition of cdk1, cells can be arrested in prophase. Our data suggest that plk1 inhibition serves as another means to stabilize the transient prophase state. The mechanism may indeed be similar by disabling one of the switches: plk1 is involved in the positive feedback loop activating cdk1, primarily acting on the cdc25C1 phosphatase (Gheghiani et al., 2017). Thus, plk1 inhibition may interfere with the positive feedback loop activating cdk1, while the PP2A:B55 branch remains intact. In this context, prolonged prophase may be an ideal condition to analyze the complex signaling network regulating mitotic entry.

In conclusion, despite advances in the field, we are still far from a complete understanding of the sequence of events leading up to mitotic entry, one of the most critical transitions in a cell’s lifetime. We need to understand how the duration of individual events is determined, how they depend on one another, and ultimately we need to identify all the molecular modifications that underlie these changes. Identification and characterization of the prolonged prophase state will greatly facilitate addressing these important questions, as it allows collection of a homogenous population of pro-phase-like cells, manipulate their mechanical environment, and profile them using cellular, molecular and biophysical assays.

## Materials and methods

### Mammalian cell culture and drug treatments

HeLa Kyoto-WT and U2-OS cell lines were kind gifts of the Ellenberg group (EMBL, Heidelberg, Germany), HeLa Kyoto EB3-mEGFP-mCherry-CenpA and HeLa Kyoto H2B-mCherry-mEGFP-PCNA were kindly provided by Daniel Gerlich (IMBA, Vienna, Austria). HeLa Kyoto LifeAct-mEGFP cells were gifted by Timo Betz (University of Göttingen, Göttingen, Germany), and hTERT RPE-1 cells by Luis Pardo (MPI-NAT, Göttingen, Germany). HeLa Kyoto Cyclin B1-mVenus was a kind gift from Jonathan Pines (ICR, London, England). HeLa Kyoto IBB-EGFP and HeLa Kyoto Eevee-spCDK-FRET (Sugiyama et al., 2023) were generated using standard procedures by random integration. HeLa-Kyoto-2xZFN-mEGFP-Nup107 #26 was purchased from CLS Cell Lines Service GmbH. All HeLa and U2-OS cells were cultured in Dulbecco’s Modified Eagle Medium (DMEM) supplemented with 10% Fetal Bovine Serum (FBS), 1% of 100 mM Sodium Pyruvate, and 1% of 200 mM L-Glutamine, all from Gibco, Thermo Fisher Scientific. hTERT RPE-1 cells were cultured in DMEM/F-12, GlutaMAX™ Supplement (Gibco, Thermo Fisher Scientific) supplemented with 10% FBS and 10 µg/ml hygromycin B.

For synchronization, cells were treated with 2 mM (HeLa, U2-OS) or 5 mM (hTERT RPE-1) thymidine in DMEM. The treatment was conducted either after at least four hours from seeding, after ensuring the cells were properly attached, or after 24 hours. 24 hours after the thymidine treatment, the medium was removed and fresh medium containing either Dimethylsulfoxide (DMSO), BI 2536 (MedChemExpress, pre-dissolved in DMSO), GSK461364 (MedChemExpress, pre-dissolved in DMSO), or RO 3306 (Sigma-Aldrich, pre-dissolved in DMSO) was added.

### Live-cell imaging experiments and dyes

Cells were imaged in FluoroBrite DMEM (Gibco, Thermo Fisher Scientific) supplemented as above. Typically, 10,000 or 12,000 cells were seeded in 96-well plates with cover glass bottom (Zell Kontakt). Dyes, such as 100 nM 5-SiR-Hoechst or 5-580CP-Hoechst (kind gifts from the Lukinavičius group, MPI-NAT in Göttingen, (Lukinavičius et al., 2015; Bucevičius et al., 2019)) to label DNA, was added to the medium prior imaging.

Imaging was done on either a Visitron Systems spinning disk confocal microscope based on a Nikon Ti2 microscope body and a Hamamatsu CSU-W1 scan head equipped with an OkoLab incubator set to 37°C with 5% CO_2_ with an airflow of 0.3 l/minute, or a Zeiss LSM 880 laser scanning confocal microscope equipped with a Pecon incubator box set to 37°C with 5% CO_2_. A Nikon Plan Apo VC, 20x Air objective (NA 0.75), Nikon Plan Apo, 40x Air objective (NA 0.95), Nikon Plan Apo TIRF AC, 100x Oil objective (NA 1.49) or Zeiss Plan Apo 40x Air (NA 0.95) objective lens were used for imaging. The 458, 488, 514, 561 and 640 nm excitation lasers were used dependent on the experimental setup, with a frequency of imaging each 5 or 10 minutes with 2 z planes 5 µm apart. Exposure times and laser powers were optimized in prior experiments to ensure that imaging does not interfere with cell survival and normal division timing.

### Immunofluorescence experiments and antibodies

For immunofluorescence experiments cells were seeded in DMEM without phenol red with supplements as above. Either 60,000 or 90,000 cells were seeded on #1.5 18 mm round cover slips (Paul Marienfield EN) in a 12-well plate. Fixation was conducted using 3% formaldehyde solution (Electron Microscopy Sciences) in PHEM buffer (1.8% PIPES, 4.2% HEPES, 0.38% EGTA, and 0.05% MgSO_4_.7H2O, adjusted to pH 7.0 with KOH) for 10 minutes. Cells were then permeabilized using 0.5% Triton X-100 in 1xPBS for 10 minutes and subsequently washed twice with 1xPBS. For storage overnight, the cover slips were stored in 1xPBS solution containing 0.02% Sodium Azide. The next day, autofluorescence was quenched using 100 mM ammonium-chloride and 100 mM glycine in 1xPBS for 20 minutes. The cover slips were then washed with 0.1% Triton X-100 in 1xPBS and blocked using 3% BSA (Roth) in 0.1% Triton X-100 in 1xPBS for 30 minutes. Primary antibodies were incubated for 1 hour using the following antibodies: 1:500 rabbit anti-H3S10ph (Sigma-Aldrich, Merck, 06-570), 1:500 mouse anti-cyclin B1 (BD Pharmingen, 554177), and mouse anti-phospho-Histone H2A.X (Ser139) (Sigma Aldrich, Merck, 05-636). Next, coverslips were washed again with 0.1% Triton X-100 in 1xPBS before and incubated with the secondary antibodies for another 1 hour in the dark. The secondary antibodies included Alexa Fluor 594 Alpaca Anti-Rabbit IgG (Jackson ImmunoResearch, 611-585-215), Alexa Fluor 594 Alpaca Anti-Mouse IgG (Jackson ImmunoResearch, 615-585-215), and Alexa Fluor 488 Alpaca Anti-Mouse IgG (Jackson ImmunoResearch, 615-545-214). 2 µg/ml Hoechst 33342 (Santa Cruz Biotechnology) was supplied with the secondary antibody for staining DNA. Following two additional washes with 0.1% Triton X-100 in 1xPBS and then 1xPBS, coverslips were mounted on SuperFrost slides using ProLong Diamond mounting medium (Invitrogen) and sealed with nail polish. Immunofluorescence samples were imaged on the same spinning disk microscope described above using the 40x Air or 100x Oil objective.

### Image analysis

ND files (VisiView 5.0, Visitron Systems) were opened using the Bio-Formats plugin by Open Microscopy Environment (OME) in Fiji/ImageJ (Schindelin et al., 2012). The z planes were typically maximum intensity projected. To produce a label image, movies were downsampled by half, denoised with Gaussian blurring (sigma of 3), and used as input to the StarDist plugin (Schmidt Uweand Weigert, 2018; Weigert et al., 2020) with the fluorescent nuclei model. Later, label images were tracked using the Trackmate plugin in Fiji (Tinevez et al., 2017; Ershov et al., 2022).

For quantifying mitotic entry, the number of nuclei in the region were counted on the first frame, then with each NEBD event, the number of cells that have undergone NEBD was counted and the cumulative percentage was calculated.

For measuring the number of gH2ax foci, the Find Maxima function of ImageJ was used with a prominence >100, and all spots detected within each nuclei were counted.

For quantifying chromosome condensation, images and labels were used as input to Knime (Berthold et al., 2007). In the Knime workflow, labels smaller than 1,000 pixels and segments touching the borders were removed. After renaming labels to match each nucleus, several features were calculated for each nucleus, including the standard deviation of fluorescence intensity and the mean fluorescence intensity. The global background was subtracted from the mean intensity of each nucleus at all time frames, and then the standard deviation was divided by the outcome to correct for bleaching.

For measuring IBB, LBR, and Cyclin B1 nuclear and cytoplasmic intensities, the same Knime workflow was used with modifications. For generating cytoplasmic labels, nuclear labels were dilated by 10 pixels using the Morphological Labeling Operations node, and then a cytoplasmic “ring” label was produced using the Voronoi Segmentation and by subtracting the nuclei label from the 10-pixel dilated-nuclei label. For IBB and LBR, the mean of the cytoplasm subtracted by the global background was normalized to the mean of the fluorescence intensity in the cytoplasm in the first 5 frames.

For measuring Eevee-spCDK biosensor FRET, the same Knime workflow was used with some modifications, where the YFP and CFP mean fluorescence intensities were measured, background subtracted, and divided by each other at every time frame for each nucleus.

For quantifying microtubule dynamics using EB3 comets, we used the uTrack software (Jaqaman et al., 2008) and its PlusTipTracker package (Applegate et al., 2011) in MatLab (MathWorks Inc., Version 9.11 (R2021b)). The 2D microtubule plus-ends object tracking was used for comet detection with a maximum of 4 frames gap-closing.

For quantifying centroid position, cells were imaged over time with 8 Z-steps and 1 µm step size. Single cells were cropped, and from the maximum Z-projection of their movies, we created label images and tracked the labels over time using the TrackMate plugin (Tinevez et al., 2017; Ershov et al., 2022). Using a macro, we then measured the centroid position, where the integrated density of the area inside the mask was measured in all Z-positions, multiplied by the number of Z-step, and the sum of all the products of all Z-steps was divided by the sum of the integrated densities from the Z-steps. The centroid positions over time were normalized to the average position in the first 5 frames of imaging.

For quantifying LifeAct-GFP signal, masks of LifeAct signal were created by Gaussian blurring (sigma 3) and later thresholding using the Huang method (Huang and Wang, 1995). The holes in the masks were filled and the watershed method was used to mark the borders of the LifeAct signal. At last, using the previously described workflow in Knime, the eccentricity was measured inside of these masks around the time of NEBD and normalized to the mean of these values in the 10 minutes following NEBD.

All mathematical operations applied on the data and data alignment and organization were conducted using Excel (Microsoft), plots and statistical tests were produced using GraphPad Prism (GraphPad Software) or the R statistical programming language. Figures were created using Affinity Publisher.

### Phosphoproteomics

Four different samples of HeLa cells as shown in Figure 5 were collected, each in three biological replicates. Cells were pelleted and lysed in SDS lysis buffer (4% [w/v] SDS, 150 mM Hepes-NaOH pH 7.5, 1x Halt™ Protease-Inhibitor-Cocktail (Thermo Scientific)) by incubation for 5 min at 99°C followed by sonication (alternating 30 s on- and 30 s off-cycles at highest intensity output (Bioruptor, Diagenode)); and by another incubation for 5 min at 99°C.

Protein concentrations were determined using Pierce™ BCA Protein Assay Kit - Reducing Agent Compatible (Thermo Scientific). Proteins were reduced with 10 mM dithiothreitol (DTT) for 30 min at 37°C, 300 rpm, and alkylated with 40 mM iodoacetamide (IAA) for 30 min at 25°C, 300 rpm, in the dark. The reaction was quenched with 10 mM DTT for 5 min at 25°C, 300 rpm. Aliquots containing 0.4 mg of protein for each replicate of each sample were further processed. Samples were diluted to final 0.4% [w/v] SDS and 1 mM MgCl_2_ was added. DNA content was digested using 1250U of Pierce™ Universal Nuclease (Thermo Scientific) for 2 hrs at 37°C, 300 rpm. For further processing, aliquots of 240 ug per sample were taken and all samples were pooled in equal protein amounts to serve as reference samples in TMT multi batch normalisations.

Separate and pooled samples were cleaned up by single-pot, solid-phase-enhanced sample preparation protocol by Hughes et al., 2019. Briefly, carboxylate modified magnetic beads (Cytiva) were added at a 1:10 protein-to-bead mass ratio and an equal volume of 100% EtOH was added. Beads were washed three times with 80% [v/v] EtOH. Proteins were digested in 50 mM TEAB containing trypsin (Promega) at a 1:20 enzyme-to-protein ratio overnight at 37 °C, 1000 rpm. Peptides were recovered from the magnetic beads and beads were washed once with 100 µl water. Pooled samples were split into six individual reference samples and further processed separately. Samples were dried in a speed vac concentrator to minimal volume (approx. 10 µl). TMTsixplex™ (Thermo Scientific) labeling was performed using three labeling batches according to manufacturer’s instructions. Briefly, 41 µl of 100% acetonitrile (ACN) were added to dried labels and dissolved by occasional vortexing. Labeling reagents were transferred to sample vials, following incubation at room temperature, 1 h; and quenched with 8 µl of 5% [v/v] hydroxylamine for 15 min at room temperature. Samples were combined including pooled reference channels. 1/20 of pooled samples were separately dried in a speed vac concentrator for whole proteome analysis.

Residual samples were subjected to phosphopeptide enrichment according to the EasyPhos protocol (Humphrey et al., 2018). Briefly, sample volumes were adjusted to 800 µl with water and subsequently to final 228 mM KCl, 3.9 mM KH_2_PO_4_, 38% [v/v] ACN, 4.5% [v/v] trifluoroacetic acid (TFA). 15 mg TiO_2_ beads (GL Sciences) in 80% [v/v] ACN, 6% [v/v] TFA were added (beads-to-protein ratio of ˃10:1). Samples were incubated for 20 min at 40°C, 1000 rpm; and washed in 60% [v/v] ACN, 1% [v/v] TFA four times and finally resuspended in 80% [v/v] ACN, 0.5% [v/v] acetic acid and mounted on top of empty columns (Harvard Apparatus). Phosphopeptides were eluted in two steps with 40% [v/v] ACN, 15% [v/v] NH_4_OH. Samples were snap-frozen in liquid nitrogen.

Whole proteome and phosphopeptide samples were cleaned-up using C18 MicroSpin columns (Harvard Apparatus) according to manufacturer’s instructions. Briefly, columns were equilibrated using 100% ACN; 50% [v/v] ACN, 0.1% [v/v] formic acid (FA); 0.1% [v/v] FA. Samples were loaded twice, washed three times with 0.1% [v/v] FA, and eluted twice with 50% [v/v] ACN, 0.1% [v/v] FA and once with 80% [v/v] ACN, 0.1% [v/v] FA. Eluants were dried in a speed vac concentrator and dissolved in 10 mM NH_4_OH pH 10, 5% [v/v] ACN for separation on an Xbridge C18 column (Waters) using an Agilent 1100 series chromatography system. The column was operated at a flow rate of 60 µl/min with a buffer system of buffer A (10 mM NH_4_OH pH 10) and buffer B (10 mM NH_4_OH pH10, 80% [v/v] ACN). Peptide separation was performed over 64 min using the following gradient: 5% B (0-7 min), 8-30% B (8-42 min), 30-50% B (43-50 min), 90-95% B (51-56 min), 5% B (57-64 min). The first 6 min were collected as flow-through (FT), followed by 48 x 1 min fractions, which were reduced to 12 fractions by concatenated pooling. Fractionated whole proteome and phosphopeptide samples were dried in a speed vac concentrator and subjected to LC-MS/MS analysis.

### LC-MS/MS analysis

Dried peptide samples were dissolved in 2% [v/v] ACN, 0.1% [v/v] TFA and injected onto a C18 Pep-Map100-trapping column (Thermo Scientific, 0.3 x 5 mm, 5 μm) connected to an in-house packed C18 analytical column (Dr Maisch GmbH, 75 μm x 300 mm). Columns were equilibrated using 98% buffer A (0.1% [v/v] FA), 2% buffer B (80% [v/v] ACN, 0.1% [v/v] FA). Liquid chromatography was performed using an UltiMate-3000 RSLC nanosystem (Thermo Scientific). Phosphopeptides were analysed for 60 min using a two-step linear gradient (5% to 34% buffer B in 34 min; 34% to 50% buffer B in 10 min) followed by a 5 min washing step at 90% of buffer B. Whole proteome peptides were analysed for 120 min, and a buffer B linear gradient of 7% to 45% over 105 min was applied followed by washing at 90% buffer B for 5 min. Eluting peptides were analysed using an Orbitrap Fusion Lumos Tribrid Mass Spectrometer (Thermo Scientific). A synchronous precursor selection (SPS) MS3 method was used with the following MS settings for phosphopeptides: MS1 scan range, 350–2000 m/z; MS1 resolution, 120,000 FWHM; AGC target MS1, 5E5; maximum injection time MS1, 50 ms; collision energy 1, 35%; charge states, 2+ to 7+; dynamic exclusion, 20 s; MS2 resolution, 30,000 FWHM, AGC target MS2, 5e5; maximum injection time MS2, 90 ms; fixed first mass MS2, m/z 132; MS3 resolution, 60,000 FWHM; collision energy 2, 45%; number of notches, 10; AGC target MS3, 5e5; maximum injection time MS3, 118 ms; fixed first mass MS3, m/z 120. For the analysis of whole proteome peptides, the following MS settings were used: MS1 scan range, 350–2000 m/z; MS1 resolution, 120,000 FWHM; AGC target MS1, 5E5; maximum injection time MS1, 50 ms; collision energy 1, 38%; charge states, 2+ to 7+; dynamic exclusion, 40 s; MS2 resolution, 15,000 FWHM, AGC target MS2, 2.5e5; maximum injection time MS2, 40 ms; fixed first mass MS2, m/z 132; MS3 resolution, 60,000 FWHM; collision energy, 45%; number of notches, 10; AGC target MS3, 2.5e5; maximum injection time MS3, 118 ms; fixed first mass MS3, m/z 120.

### Phosphoproteomics Data Analysis

MS raw files were processed using MaxQuant version 2.0.3.0 (Cox and Mann, 2008) with default settings except for: fixed modification, carbamidomethylation (C); variable modifications, oxidation (M), acetylation (N-term), phosphorylation (S,T,Y) (only for phosphopeptide data). MS3 reporter ions (TMTsixplex™) were selected as quantification type, defining pooled sample channels as reference channels. Human canonical protein sequences (UP000005640) were downloaded from UniProt knowledgebase (Reviewed sequences, Swiss-Prot; date of download: 11.07.2022; number of protein sequences: 20,598).

Following steps in data analysis were conducted in Perseus (Tyanova et al., 2016). For each phosphorylation site, a leading protein was selected based on the list of potential candidate proteins for this site reported by MaxQuant. The official gene name and Uniprot accession of the leading protein were used in the subsequent analyses. Reporter ion intensities for each phosphorylation site identified at one of the three multiplicity levels were extracted from MaxQuant “Phospho(S, T, Y).txt’’ output table. Potential contaminants, reversed sequences, and phosphorylation sites identified with localization probability <0.75 as determined by MaxQuant were excluded from further analysis. If a phosphorylation event contained <70% nonzero (valid) intensity values, the event was considered as not quantified in this labeling experiment. Otherwise, missing values were imputed for each TMT channel/replicate individually by random sampling from a Gaussian distribution with a width of 0.3 and a downshift of 1.8 of the log2-transformed intensities. Log2-transformed reporter ion intensities were then normalized using Tukey median polishing individually for each labeling experiment in R statistical programming language and later subjected to statistical testing using the Limma package (Ritchie et al., 2015).

Linear models were fitted using least squares method from limma R library. P-values were estimated with the empirical Bayes method and multiple hypothesis testing correction was done with the Benjamini and Hochberg method. A significance cutoff of 0.05 adjusted p-value was used.

For the comparison shown in Fig. 5D, E and Fig. 6, treatment-contrast parameterization was established using a 2-by-2 factorial design for the linear modeling, with the following resulting comparison [BI2536.attached - BI2536.suspended] - [DMSO.attached - DMSO.suspended] (Fig.5B).

For figure S4C & D, all the phosphopeptides tested for differential expression, including insignificant hits, were converted to the asterisk format, where the phosphorylated amino acid is marked with an asterisk (*) after the phospho-acceptor (S, T, Y). Peptide sequences, log-fold changes and p.values of differential expression were submitted for kinase prediction at phosphosite.org (Hornbeck et al., 2015) KinaseLibraryAction functional enrichment analysis based on differential expression. A p-value threshold of 0.05 was used to discriminate between foreground and the background sets.

Other R packages were also used to create the graphs shown in Figure 5 and 6, including ggplot2 (Wickham, 2009), GO.db, GOstats, and BioConductor (Huber et al., 2015).

## Supplementary figure legends

**Figure S1.**
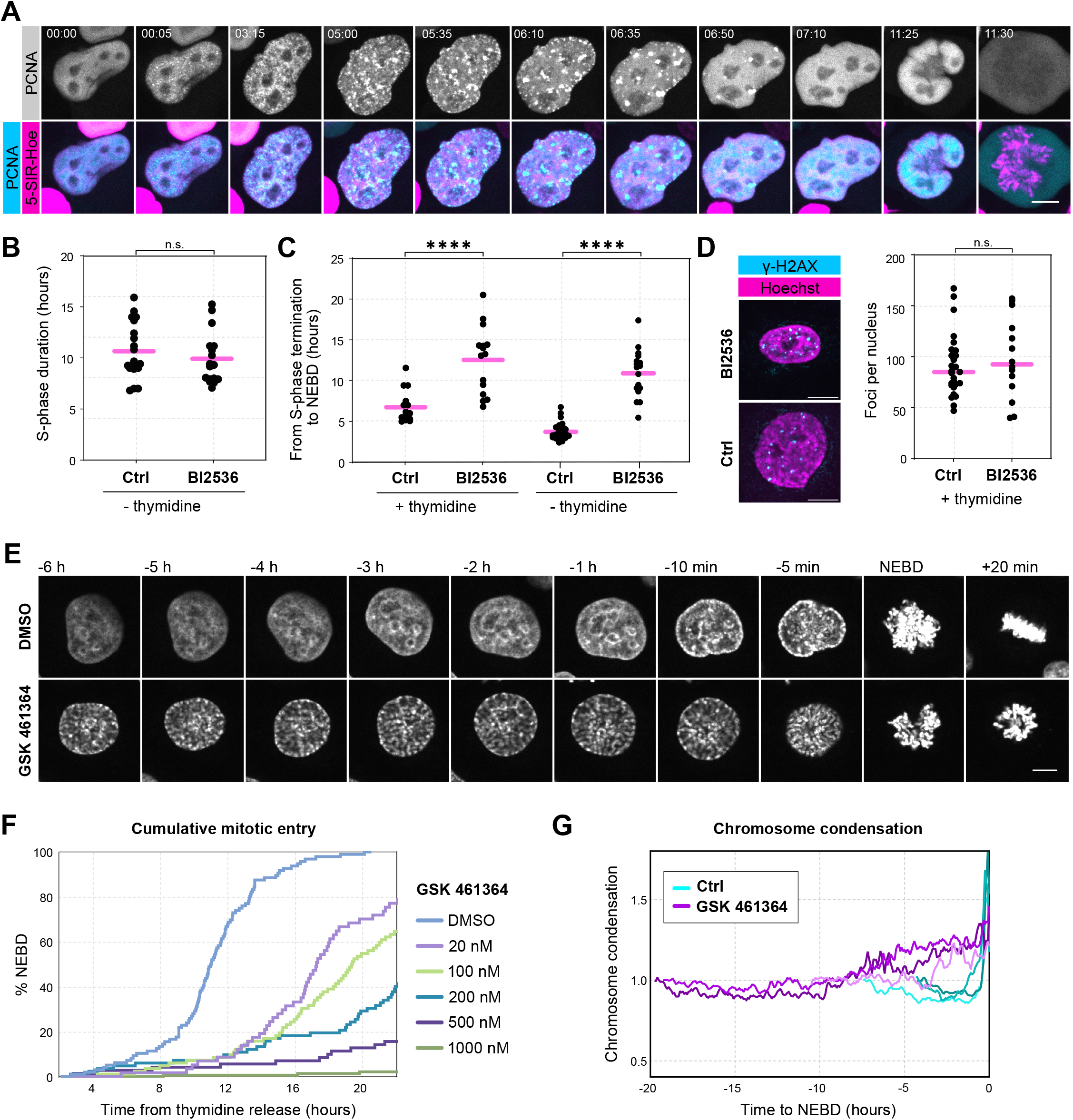
**(A)** HeLa cells expressing PCNA-mEGFP revealing S-phase progression after thymidine release. **(B)** Quantification of the S-phase duration in control and BI2536-treated cells. **(C)** Quantification of the duration from S-phase termination to NEBD and with and without thymidine synchronization in HeLa Kyoto cells measured by PCNA labeling. **** corresponds to p < 0.0001 and ns corresponds to p > 0.05 (p = 0.4028) **(D)** Immunofluorescence images of HeLa Kyoto cells stained for γ-H2AX antibodies. Example images (left) and quantification of the number of g-H2AX foci per nuclei (right). The control and treated cells were analyzed 8 and 17 h after thymidine release, respectively. ns corresponds to p > 0.05 (p = 0.5467). **(E)** Montages of HeLa cells showing mitotic entry of exemplary control and GSK 461364-treated cells stained with 5-SiR-Hoechst. **(F)** Cumulative plots of NEBD timing in response to increasing concentrations of GSK 461364 in HeLa Kyoto cells. **(G)** Quantification of chromosome condensation by standard deviation of fluorescence intensity of individual HeLa Kyoto cells on recording similar to shown in (E). For (B), (C), and (D): significance testing was done using unpaired Student’s *t*-test with Welch’s correction. For (A), (D), and (E): scale bars are 10 µm.

**Figure S2.**
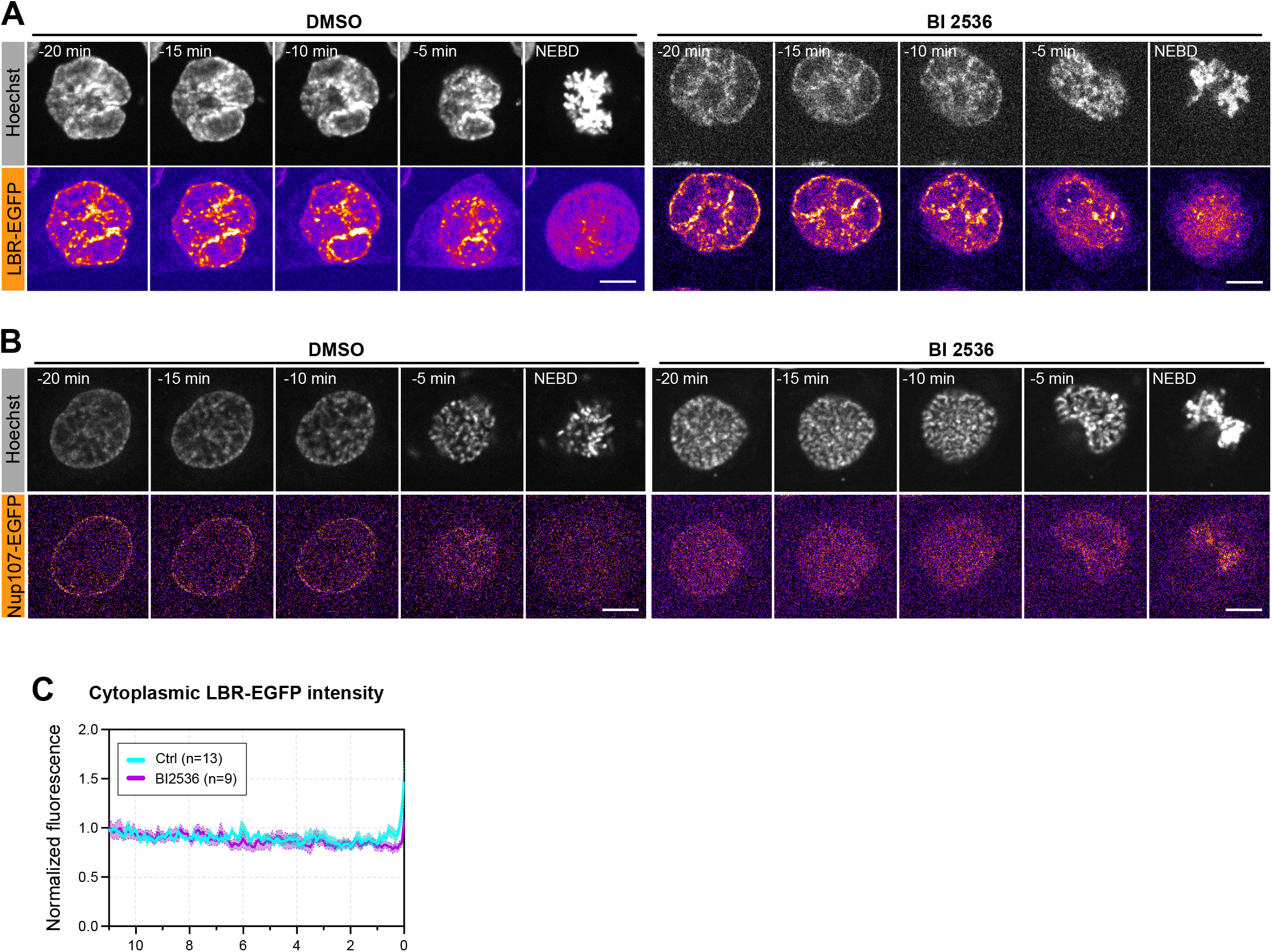
**(A)** Selected frames from a time lapse showing the localization of LBR-EGFP in the time leading to NEBD in control and BI 2536-treated cells. **(B)** Selected frames from a time lapse showing the localization of Nup107-EGFP in the time leading to NEBD in control and BI 2536-treated cells. **(C)** Quantification of LBR-EGFP mean cytoplasmic fluorescence intensity in control and BI 536-treated cells on recordings similar to (A). Data is normalized to the mean intensity in the first 5 frames of imaging.

**Figure S3.**
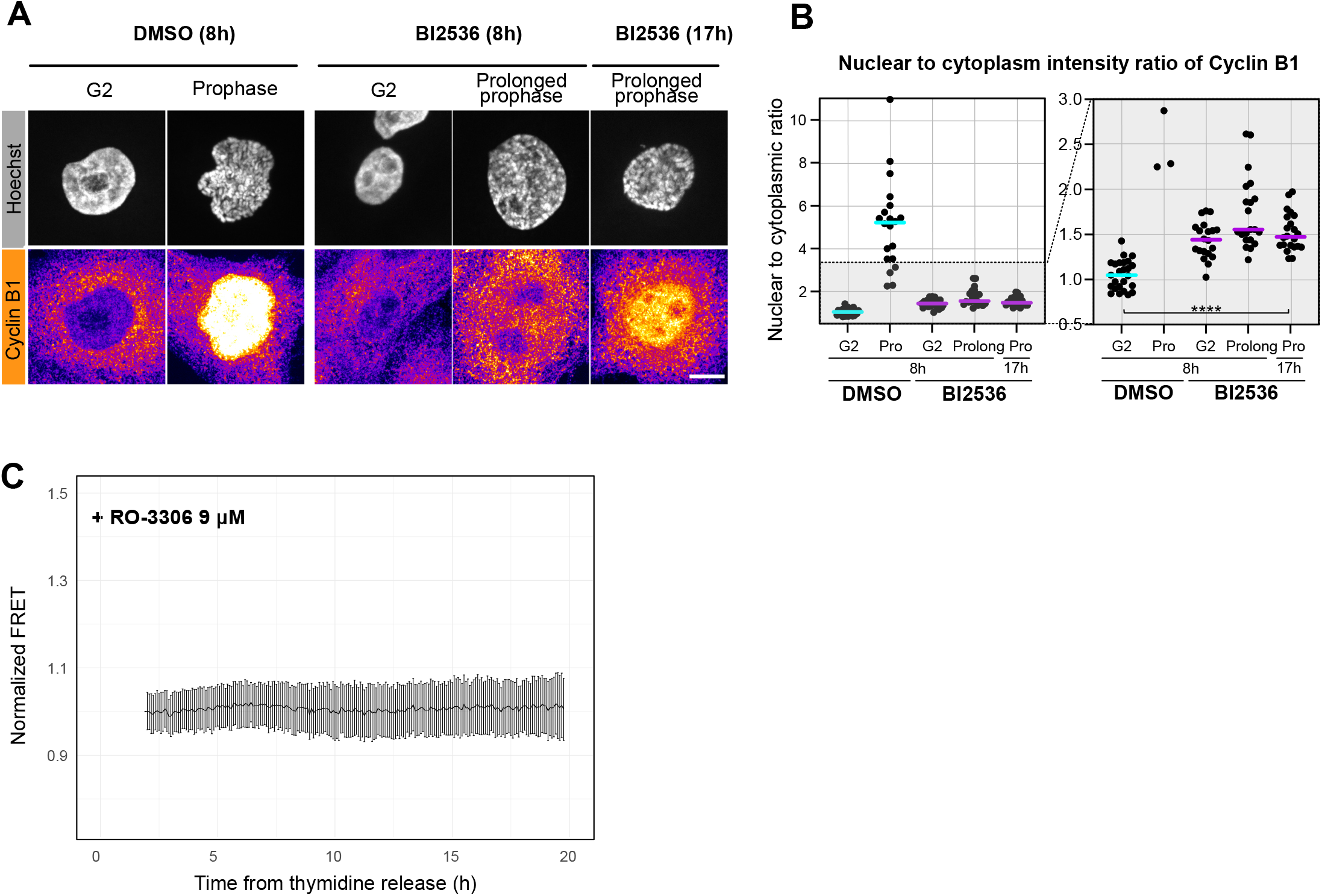
**(A)** Immunofluorescence of cyclin B1 in control and BI 2536-treated cells fixed at different time points after thymidine release. Scale bar is 10 µm. **(B)** Quantification of nuclear to cytoplasmic ratio of fluorescence intensity of cyclin B1 in control and treated cells on images similar to (A). Inset magnifying the lower range is shown on the right panel. **(C)** Quantification of FRET ratio changes of the Eevee-spCDK FRET probe similar to that shown in Fig. 4C in RO 3306-treated population of cells.

**Figure S4.**
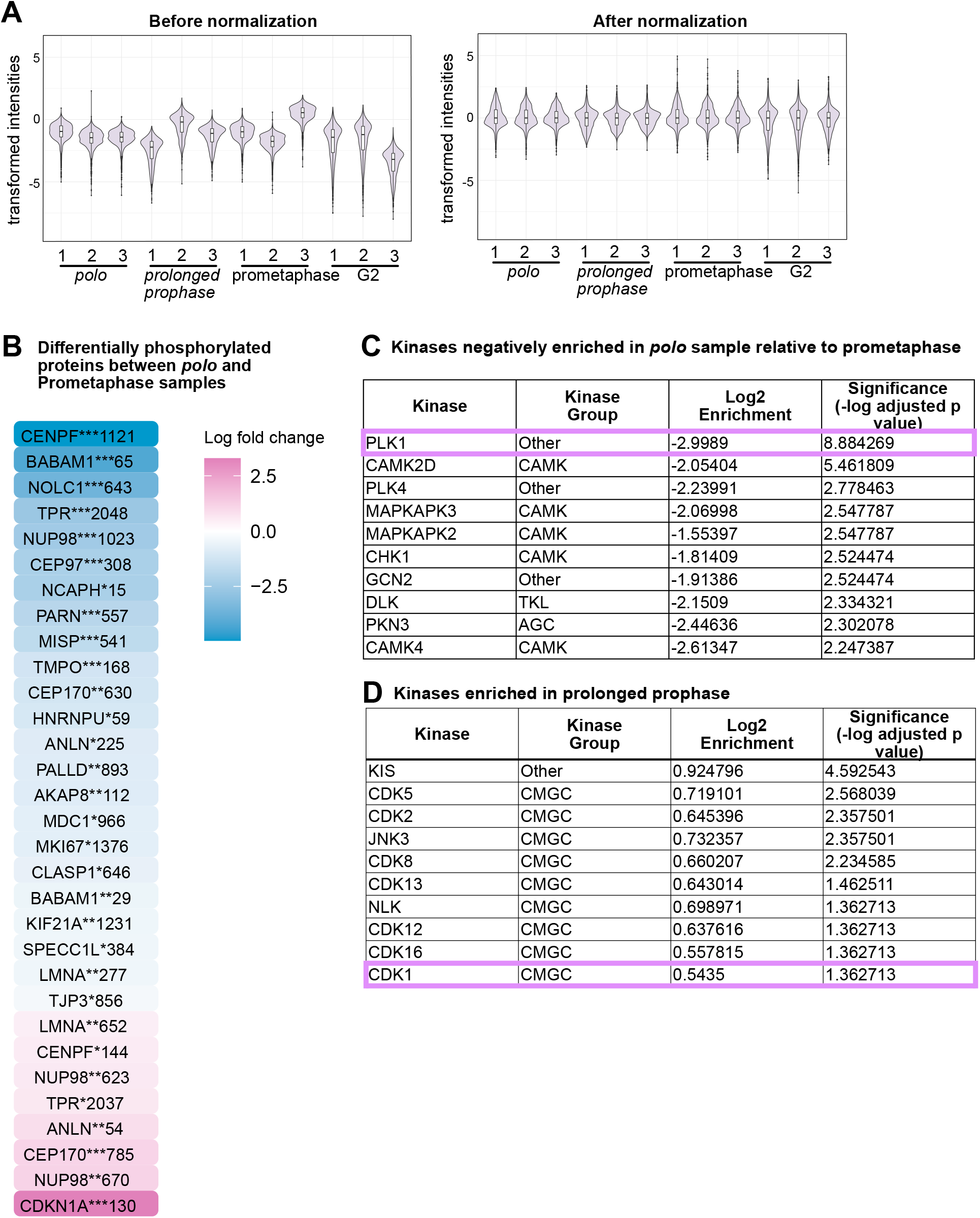
**(A)** Violin plots showing the distribution of the log(2) transformed intensities of the phosphosites detected in the 12 samples (4 samples in 3 replicates) before and after normalization. **(B)** Differentially phosphorylated peptides in *polo* cells in comparison to prometaphase cells that were detected using linear modeling. The gene names of the corresponding peptides are show, the asterisks mark the adjusted p-value with ***, **, * corresponding to p ≤ 0.001, 0.01, and 0.05, respectively, followed by the position of the phosphosite within the amino acid sequence of the protein. The color coding corresponds to the log(2) fold change of intensity values. **(C)** List of kinases least enriched in the polo sample in comparison to the prometaphase sample after applying linear modeling. **(D)** List of kinases most enriched in the prolonged prophase sample after applying linear modeling.

## Acknowledgements

We would like to thank Jan Ellenberg (EMBL, Heidelberg, Germany), Daniel Gerlich (IMBA, Vienna, Austria), Luis Pardo (MPI-NAT), Jonathan Pines (Institute of Cancer Research, London, England) and Timo Betz (University of Göttingen) for providing cell lines, as well as Grazvydas Lukinavičius (MPI-NAT) and Kazuhiro Aoki (NIBB, Okazaki, Japan) for reagents. We would like to thank Juliane Liepe (MPI-NAT), Arshad Desai (Ludvig Cancer Research, UCSD, USA), Matthias Dobbelstein (Molecular Oncology, UMG, Göttingen, Germany) and Jonathon Pines (Institute of Cancer Research, London, UK) for comments on the manuscript. We would like to acknowledge the support of MPI-NAT’s core facilities, the Proteomics core facility in particular. Research in P.L.’s group is funded by the Max Planck Society, M.G. is enrolled in the IMPRS Molecular Biology graduate programme.

## References

Ali, A., C. Vineethakumari, C. Lacasa, and J. Lüders. 2023. Microtubule nucleation and γTuRC centrosome localization in interphase cells require ch-TOG. Nat. Commun. 14:289. doi:10.1038/s41467-023-35955-w.

Applegate, K.T., S. Besson, A. Matov, M.H. Bagonis, K. Jaqaman, and G. Danuser. 2011. plusTipTracker: Quantitative image analysis software for the measurement of microtubule dynamics. J. Struct. Biol. 176:168–184. doi:10.1016/j.jsb.2011.07.009.

Beaudouin, J., D. Gerlich, N. Daigle, R. Eils, and J. Ellenberg. 2002. Nuclear envelope breakdown proceeds by microtubule-induced tearing of the lamina. Cell. 108:83–96. doi:10.1016/s0092-8674(01)00627-4.

Bucevičius, J., J. Keller-Findeisen, T. Gilat, S.W. Hell, and G. Lukinavičius. 2019. Rhodamine–Hoechst positional isomers for highly efficient staining of heterochromatin. Chem. Sci. 10:1962–1970. doi:10.1039/C8SC05082A.

Bucevicius, J., G. Kostiuk, R. Gerasimaite, T. Gilat, and G. Lukinavicius. 2020. Enhancing biocompatibility of rhodamine fluorescent probes by a neighbouring group effect. Chem. Sci. 11. doi:10.1039/D0SC02154G.

Cassimeris, L. 2002. The oncoprotein 18/stathmin family of microtubule destabilizers. Curr. Opin. Cell Biol. 14:18–24. doi:10.1016/S0955-0674(01)00289-7.

Chen, N.-P., J. Aretz, and R. Fässler. 2022. CDK1–cyclin-B1-induced kindlin degradation drives focal adhesion disassembly at mitotic entry. Nat. Cell Biol. 24:723–736. doi:10.1038/s41556-022-00886-z.

Conduit, P.T., A. Wainman, and J.W. Raff. 2015. Centrosome function and assembly in animal cells. Nat. Rev. Mol. Cell Biol. 16:611–624. doi:10.1038/nrm4062.

Cox, J., and M. Mann. 2008. MaxQuant enables high peptide identification rates, individualized p.p.b.- range mass accuracies and proteome-wide protein quantification. Nat. Biotechnol. 26:1367– 1372. doi:10.1038/nbt.1511.

Crncec, A., and H. Hochegger. 2019. Triggering mitosis. FEBS Lett. 593:2868–2888. doi:10.1002/1873-3468.13635.

Cuylen, S., C. Blaukopf, A.Z. Politi, T. Müller-Reichert, B. Neumann, I. Poser, J. Ellenberg, A.A. Hyman, and D.W. Gerlich. 2016. Ki-67 acts as a biological surfactant to disperse mitotic chromosomes. Nature. 535:308–312. doi:10.1038/nature18610.

Daub, H., J.V. Olsen, M. Bairlein, F. Gnad, F.S. Oppermann, R. Körner, Z. Greff, G. Kéri, O. Stemmann, and M. Mann. 2008. Kinase-Selective Enrichment Enables Quantitative Phosphoproteomics of the Kinome across the Cell Cycle. Mol. Cell. 31:438–448. doi:10.1016/j.molcel.2008.07.007.

Dephoure, N., C. Zhou, J. Villén, S.A. Beausoleil, C.E. Bakalarski, S.J. Elledge, and S.P. Gygi. 2008. A quantitative atlas of mitotic phosphorylation. Proc. Natl. Acad. Sci. 105:10762–10767. doi:10.1073/pnas.0805139105.

Dultz, E., E. Zanin, C. Wurzenberger, M. Braun, G. Rabut, L. Sironi, and J. Ellenberg. 2008. Systematic kinetic analysis of mitotic dis- and reassembly of the nuclear pore in living cells. J. Cell Biol. 180:857–865. doi:10.1083/jcb.200707026.

Ershov, D., M.-S. Phan, J.W. Pylvänäinen, S.U. Rigaud, L. Le Blanc, A. Charles-Orszag, J.R.W. Conway, R.F. Laine, N.H. Roy, D. Bonazzi, G. Duménil, G. Jacquemet, and J.-Y. Tinevez. 2022. TrackMate 7: integrating state-of-the-art segmentation algorithms into tracking pipelines. Nat. Methods. 19:829–832. doi:10.1038/s41592-022-01507-1.

Farina, F., N. Ramkumar, L. Brown, D. Samandar Eweis, J. Anstatt, T. Waring, J. Bithell, G. Scita, M. Thery, L. Blanchoin, T. Zech, and B. Baum. 2019. Local actin nucleation tunes centrosomal microtubule nucleation during passage through mitosis. EMBO J. 38:e99843. doi:10.15252/embj.201899843.

Ferenz, N.P., and P. Wadsworth. 2007. Prophase Microtubule Arrays Undergo Flux-like Behavior in Mammalian Cells. Mol. Biol. Cell. 18:3993–4002. doi:10.1091/mbc.E07-05-0420.

Gavet, O., and J. Pines. 2010. Activation of cyclin B1-Cdk1 synchronizes events in the nucleus and the cytoplasm at mitosis. J. Cell Biol. 189:247–259. doi:10.1083/jcb.200909144.

Gheghiani, L., D. Loew, B. Lombard, J. Mansfeld, and O. Gavet. 2017. PLK1 Activation in Late G2 Sets Up Commitment to Mitosis. Cell Rep. 19:2060–2073. doi:10.1016/j.celrep.2017.05.031.

Gilmartin, A.G., M.R. Bleam, M.C. Richter, S.G. Erskine, R.G. Kruger, L. Madden, D.F. Hassler, G.K. Smith, R.R. Gontarek, M.P. Courtney, D. Sutton, M.A. Diamond, J.R. Jackson, and S.G. Laquerre. 2009. Distinct concentration-dependent effects of the polo-like kinase 1-specific inhibitor GSK461364A, including differential effect on apoptosis. Cancer Res. 69:6969–6977. doi:10.1158/0008-5472.CAN-09-0945.

Hirota, T., D. Gerlich, B. Koch, J. Ellenberg, and J.-M. Peters. 2004. Distinct functions of condensin I and II in mitotic chromosome assembly. J. Cell Sci. 117:6435–6445. doi:10.1242/jcs.01604.

Hirota, T., J.J. Lipp, B.-H. Toh, and J.-M. Peters. 2005. Histone H3 serine 10 phosphorylation by Aurora B causes HP1 dissociation from heterochromatin. Nature. 438:1176–1180. doi:10.1038/nature04254.

Hornbeck, P.V., B. Zhang, B. Murray, J.M. Kornhauser, V. Latham, and E. Skrzypek. 2015. PhosphoSitePlus, 2014: mutations, PTMs and recalibrations. Nucleic Acids Res. 43:D512–D520. doi:10.1093/nar/gku1267.

Hsu, J.-Y., Z.-W. Sun, X. Li, M. Reuben, K. Tatchell, D.K. Bishop, J.M. Grushcow, C.J. Brame, J.A. Caldwell, D.F. Hunt, R. Lin, M.M. Smith, and C.D. Allis. 2000. Mitotic Phosphorylation of Histone H3 Is Governed by Ipl1/aurora Kinase and Glc7/PP1 Phosphatase in Budding Yeast and Nematodes. Cell. 102:279–291. doi:10.1016/S0092-8674(00)00034-9.

Hughes, C.S., S. Moggridge, T. Müller, P.H. Sorensen, G.B. Morin, and J. Krijgsveld. 2019. Single-pot, solid-phase-enhanced sample preparation for proteomics experiments. Nat. Protoc. 14:68–85. doi:10.1038/s41596-018-0082-x.

Humphrey, S.J., O. Karayel, D.E. James, and M. Mann. 2018. High-throughput and high-sensitivity phosphoproteomics with the EasyPhos platform. Nat. Protoc. 13:1897–1916. doi:10.1038/s41596-018-0014-9.

Jelluma, N., A.B. Brenkman, N.J.F. van den Broek, C.W.A. Cruijsen, M.H.J. van Osch, S.M.A. Lens, R.H. Medema, and G.J.P.L. Kops. 2008. Mps1 phosphorylates Borealin to control Aurora B activity and chromosome alignment. Cell. 132:233–246. doi:10.1016/j.cell.2007.11.046.

Kelly, V., A. al-Rawi, D. Lewis, G. Kustatscher, and T. Ly. 2021. Low Cell Number Proteomic Analysis Using In-Cell Protease Digests Reveals a Robust Signature for Cell Cycle State Classification. Mol. Cell. Proteomics MCP. 21:100169. doi:10.1016/j.mcpro.2021.100169.

Laurell, E., K. Beck, K. Krupina, G. Theerthagiri, B. Bodenmiller, P. Horvath, R. Aebersold, W. Antonin, and U. Kutay. 2011. Phosphorylation of Nup98 by multiple kinases is crucial for NPC disassembly during mitotic entry. Cell. 144:539–550. doi:10.1016/j.cell.2011.01.012.

Lee, K., and K. Rhee. 2011. PLK1 phosphorylation of pericentrin initiates centrosome maturation at the onset of mitosis. J. Cell Biol. 195:1093–1101. doi:10.1083/jcb.201106093.

Lénárt, P., and J. Ellenberg. 2006. Monitoring the permeability of the nuclear envelope during the cell cycle. Methods San Diego Calif. 38:17–24. doi:10.1016/j.ymeth.2005.07.010.

Lénárt, P., M. Petronczki, M. Steegmaier, B. Di Fiore, J.J. Lipp, M. Hoffmann, W.J. Rettig, N. Kraut, and J.-M. Peters. 2007. The small-molecule inhibitor BI 2536 reveals novel insights into mitotic roles of polo-like kinase 1. Curr. Biol. CB. 17:304–315. doi:10.1016/j.cub.2006.12.046.

Linder, M.I., M. Köhler, P. Boersema, M. Weberruss, C. Wandke, J. Marino, C. Ashiono, P. Picotti, W. Antonin, and U. Kutay. 2017. Mitotic Disassembly of Nuclear Pore Complexes Involves CDK1- and PLK1-Mediated Phosphorylation of Key Interconnecting Nucleoporins. Dev. Cell. 43:141–156.e7. doi:10.1016/j.devcel.2017.08.020.

Liu, K., M. Zheng, R. Lu, J. Du, Q. Zhao, Z. Li, Y. Li, and S. Zhang. 2020. The role of CDC25C in cell cycle regulation and clinical cancer therapy: a systematic review. Cancer Cell Int. 20:213. doi:10.1186/s12935-020-01304-w.

Ly, T., A. Whigham, R. Clarke, A.J. Brenes-Murillo, B. Estes, D. Madhessian, E. Lundberg, P. Wadsworth, and A.I. Lamond. 2017. Proteomic analysis of cell cycle progression in asynchronous cultures, including mitotic subphases, using PRIMMUS. eLife. 6:e27574. doi:10.7554/eLife.27574.

Neurohr, G., and D.W. Gerlich. 2009. Assays for mitotic chromosome condensation in live yeast and mammalian cells. Chromosome Res. Int. J. Mol. Supramol. Evol. Asp. Chromosome Biol. 17:145–154. doi:10.1007/s10577-008-9010-1.

Nielsen, C.F., T. Zhang, M. Barisic, P. Kalitsis, and D.F. Hudson. 2020. Topoisomerase IIα is essential for maintenance of mitotic chromosome structure. Proc. Natl. Acad. Sci. 117:12131–12142. doi:10.1073/pnas.2001760117.

Nkoula, S.N., G. Velez-Aguilera, B. Ossareh-Nazari, L.V. Hove, C. Ayuso, V. Legros, G. Chevreux, L. Thomas, G. Seydoux, P. Askjaer, and L. Pintard. 2023. Mechanisms of Nuclear Pore Complex disassembly by the mitotic Polo-Like Kinase 1 (PLK-1) in C. elegans embryos. BioRxiv Prepr. Serv. Biol. 2023.02.21.528438. doi:10.1101/2023.02.21.528438.

Peset, I., and I. Vernos. 2008. The TACC proteins: TACC-ling microtubule dynamics and centrosome function. Trends Cell Biol. 18:379–388. doi:10.1016/j.tcb.2008.06.005.

Petronczki, M., M. Glotzer, N. Kraut, and J.-M. Peters. 2007. Polo-like kinase 1 triggers the initiation of cytokinesis in human cells by promoting recruitment of the RhoGEF Ect2 to the central spindle. Dev. Cell. 12:713–725. doi:10.1016/j.devcel.2007.03.013.

Raaijmakers, J.A., R.G.H.P. van Heesbeen, J.L. Meaders, E.F. Geers, B. Fernandez-Garcia, R.H. Medema, and M.E. Tanenbaum. 2012. Nuclear envelope-associated dynein drives prophase centrosome separation and enables Eg5-independent bipolar spindle formation. EMBO J. 31:4179–4190. doi:10.1038/emboj.2012.272.

Rata, S., M.F. Suarez Peredo Rodriguez, S. Joseph, N. Peter, F. Echegaray Iturra, F. Yang, A. Madzvamuse, J.G. Ruppert, K. Samejima, M. Platani, M. Alvarez-Fernandez, M. Malumbres, W.C. Earnshaw, B. Novak, and H. Hochegger. 2018. Two Interlinked Bistable Switches Govern Mitotic Control in Mammalian Cells. Curr. Biol. CB. 28:3824–3832.e6. doi:10.1016/j.cub.2018.09.059.

Riedl, J., A.H. Crevenna, K. Kessenbrock, J.H. Yu, D. Neukirchen, M. Bista, F. Bradke, D. Jenne, T.A. Holak, Z. Werb, M. Sixt, and R. Wedlich-Soldner. 2008. Lifeact: a versatile marker to visualize F-actin. Nat. Methods. 5:605–607. doi:10.1038/nmeth.1220.

Ritchie, M.E., B. Phipson, D. Wu, Y. Hu, C.W. Law, W. Shi, and G.K. Smyth. 2015. limma powers differential expression analyses for RNA-sequencing and microarray studies. Nucleic Acids Res. 43:e47–e47. doi:10.1093/nar/gkv007.

Rottner, K., J. Hänisch, and K.G. Campellone. 2010. WASH, WHAMM and JMY: regulation of Arp2/3 complex and beyond. Trends Cell Biol. 20:650–661. doi:10.1016/j.tcb.2010.08.014.

Samejima, I., C. Spanos, K. Samejima, J. Rappsilber, G. Kustatscher, and W.C. Earnshaw. 2022. Mapping the invisible chromatin transactions of prophase chromosome remodeling. Mol. Cell. 82:696–708.e4. doi:10.1016/j.molcel.2021.12.039.

Samora, C.P., B. Mogessie, L. Conway, J.L. Ross, A. Straube, and A.D. McAinsh. 2011. MAP4 and CLASP1 operate as a safety mechanism to maintain a stable spindle position in mitosis. Nat. Cell Biol. 13:1040–1050. doi:10.1038/ncb2297.

Santos, S.D.M., R. Wollman, T. Meyer, and J.E. Ferrell. 2012. Spatial positive feedback at the onset of mitosis. Cell. 149:1500–1513. doi:10.1016/j.cell.2012.05.028.

Schweizer, N., C. Ferrás, D.M. Kern, E. Logarinho, I.M. Cheeseman, and H. Maiato. 2013. Spindle assembly checkpoint robustness requires Tpr-mediated regulation of Mad1/Mad2 proteostasis. J. Cell Biol. 203:883–893. doi:10.1083/jcb.201309076.

Steegmaier, M., M. Hoffmann, A. Baum, P. Lénárt, M. Petronczki, M. Krssák, U. Gürtler, P. Garin-Chesa, S. Lieb, J. Quant, M. Grauert, G.R. Adolf, N. Kraut, J.-M. Peters, and W.J. Rettig. 2007. BI 2536, a potent and selective inhibitor of polo-like kinase 1, inhibits tumor growth in vivo. Curr. Biol. CB. 17:316–322. doi:10.1016/j.cub.2006.12.037.

Sugiyama, H., Y. Goto, Y. Kondo, D. Coudreuse, and K. Aoki. 2023. Live-cell imaging provides direct evidence for a threshold in CDK activity at the G2/M transition. 2023.03.26.534249. doi:10.1101/2023.03.26.534249.

Tanenbaum, M.E., and R.H. Medema. 2010. Mechanisms of Centrosome Separation and Bipolar Spindle Assembly. Dev. Cell. 19:797–806. doi:10.1016/j.devcel.2010.11.011.

Taubenberger, A.V., B. Baum, and H.K. Matthews. 2020. The Mechanics of Mitotic Cell Rounding. Front. Cell Dev. Biol. 8.

Tinevez, J.-Y., N. Perry, J. Schindelin, G.M. Hoopes, G.D. Reynolds, E. Laplantine, S.Y. Bednarek, S.L. Shorte, and K.W. Eliceiri. 2017. TrackMate: An open and extensible platform for single-particle tracking. Methods. 115:80–90. doi:10.1016/j.ymeth.2016.09.016.

Tyanova, S., T. Temu, P. Sinitcyn, A. Carlson, M.Y. Hein, T. Geiger, M. Mann, and J. Cox. 2016. The Perseus computational platform for comprehensive analysis of (prote)omics data. Nat. Methods. 13:731–740. doi:10.1038/nmeth.3901.

van der Vaart, B., C. Manatschal, I. Grigoriev, V. Olieric, S.M. Gouveia, S. Bjelic, J. Demmers, I. Vorobjev, C.C. Hoogenraad, M.O. Steinmetz, and A. Akhmanova. 2011. SLAIN2 links microtubule plus end-tracking proteins and controls microtubule growth in interphase. J. Cell Biol. 193:1083– 1099. doi:10.1083/jcb.201012179.

Vagnarelli, P., D.F. Hudson, S.A. Ribeiro, L. Trinkle-Mulcahy, J.M. Spence, F. Lai, C.J. Farr, A.I. Lamond, and W.C. Earnshaw. 2006. Condensin and Repo-Man-PP1 co-operate in the regulation of chromosome architecture during mitosis. Nat. Cell Biol. 8:1133–1142. doi:10.1038/ncb1475.

Velez-Aguilera, G., S. Nkombo Nkoula, B. Ossareh-Nazari, J. Link, D. Paouneskou, L. Van Hove, N. Joly, N. Tavernier, J.-M. Verbavatz, V. Jantsch, and L. Pintard. 2020. PLK-1 promotes the merger of the parental genome into a single nucleus by triggering lamina disassembly. eLife. 9:e59510. doi:10.7554/eLife.59510.

Vergnolle, M.A.S., and S.S. Taylor. 2007. Cenp-F links kinetochores to Ndel1/Nde1/Lis1/dynein microtubule motor complexes. Curr. Biol. CB. 17:1173–1179. doi:10.1016/j.cub.2007.05.077.

Wilkins, B.J., N.A. Rall, Y. Ostwal, T. Kruitwagen, K. Hiragami-Hamada, M. Winkler, Y. Barral, W. Fischle, and H. Neumann. 2014. A Cascade of Histone Modifications Induces Chromatin Condensation in Mitosis. Science. 343:77–80. doi:10.1126/science.1244508.

Xu, Z., H. Ogawa, P. Vagnarelli, J.H. Bergmann, D.F. Hudson, S. Ruchaud, T. Fukagawa, W.C. Earnshaw, and K. Samejima. 2009. INCENP–aurora B interactions modulate kinase activity and chromosome passenger complex localization. J. Cell Biol. 187:637–653. doi:10.1083/jcb.200906053.

